# Mosquitoes escape looming threats by actively flying with the bow-wave induced by the attacker

**DOI:** 10.1101/2023.08.28.555052

**Authors:** Antoine Cribellier, Leonardo Honfi Camilo, Pulkit Goyal, Florian T. Muijres

## Abstract

To detect and escape from a looming threat, night-flying insects must rely on other senses than vision alone. Nocturnal mosquitoes have been described to escape looming objects in the dark, but how they achieve this is still unknown. Here, we show how night-active female malaria mosquitoes escape from a rapidly looming object that simulates the defensive action of a blood-host. By combining videography-based automatic tracking with numerical simulations of the attacker-induced airflow, we first show that night-flying mosquitoes use airflow-sensing to detect the danger and trigger their escape. Secondly, by combining these data with mechanistic movement modelling, we unravelled how mosquitoes control their escape manoeuvres: they actively steer away from the danger, and passively travel with the bow-wave produced by the attacker. Our results demonstrate that night-flying mosquitoes escaping from a looming object use the object-induced airflow both to detect the danger, and as fluid medium to move with for avoiding collision. This shows that the escape strategy of flying insects is more complex than previous visually-induces escape flight studies suggest. As mosquitoes are average-sized insects, a combined airflow-induced and visual-induced escape strategy is expected to be common amongst millions of flying insect species. Also, our research helps explain the high escape performance of mosquitoes from counterflow-based odour-baited mosquito traps. It can therefore provide new insights for the development of novel trapping techniques for integrative vector management.

## Introduction

The females of hematophagous mosquitoes must interact with their hosts to get a blood meal necessary for egg development (1). This interaction is the reason why anthropophilic mosquitoes can be vectors of many deadly diseases including malaria, yellow fever, and dengue (2). To protect themselves, humans developed vector control tools such as bed nets or traps (3, 4). But arguably, the vector control technique most widely-used by both humans and other animals is to discourage or kill host-seeking mosquitoes using hand swatting or tail swishing (5–10). In addition, flying mosquitoes can be attacked by various predators such as dragonflies, bats or birds (11–13). Despite the large amount of research on host seeking behaviours (14–17), how mosquitoes respond to host defensive strategies and to predator’s attacks has been studied very little (18, 19).

To avoid a looming threat, a mosquito will rely on a combination of so-called protean insurance behaviours – exhibiting unpredictable flight paths – and escape manoeuvres (18). The relative proportion of these two strategies explaining escape performances is dependent on light intensity and species, most likely in order to adapt to available sensory information (18). Thus, nocturnal malaria mosquitoes decrease their flight predictability in the dark by flying faster. And, both night-active malaria mosquitoes and day-active yellow fewer mosquitoes exhibit fast escape manoeuvres more often with increasing light intensity. These results suggests that mosquitoes use visual cues to detect threats and trigger their escapes, but it is striking that nocturnal mosquitoes still perform escape manoeuvres in dark conditions when limited to no visual cues are present (18).

Escape manoeuvres of flying mosquitoes have been little studied, but more is known about the escape flights of hummingbirds, moths and fruit flies. While escaping, these animals usually direct their manoeuvres away from the danger (20, 21), or towards safe zones at the flank of the attacker (22). Fruit flies, dipterans like mosquitoes, follow the so-called ‘helicopter model’ when performing an escape manoeuvre (20). According to this model, the animal manoeuvres by rotating its whole body to redirect its produced aerodynamic force vector in the direction of the intended motion (23, 24). Thus, when evading looming targets, fruit flies execute banked turns by pitching their nose-up and rolling on the side opposite to the threat location (20). They do so by modulating kinematic parameters such as wingbeat frequency and amplitude. Measurements of mosquito kinematics during take-off suggest that manoeuvring mosquitoes comply to the same ‘helicopter model’ as fruit flies do (25).

All previously mentioned escape manoeuvres are fully active manoeuvres, where the animal detects the threat and then responds using an escape manoeuvre. Most of the time, the incoming threats are detected using vision alone (20, 26). However, insects flying in the dark would need to rely on other senses. Various large ground dwelling insect are able to detect the air gust produced by an attack, which then triggers their escape (27–31). But there is no similar known example amongst smaller flying insects. Praying mantis, crickets and cockroaches use sensible hairs or cerci in order to trigger responses (28, 30, 32, 33). Smaller insects could rely on their mechanoreceptors (i.e. Johnston’s organs and sensible hairs) to detect the air gust generated by the attack in order to trigger an active response (34). Alternatively, these flying insects may also be passively pushed away by the attacker-induced airflow. Indeed, most insects have low inertia, and therefore the moving air might generate a high enough aerodynamic drag force on the insect’s body to move it away from danger, without the need of an active response.

Here, we tested how night-flying anthropophilic malaria mosquitoes (*Anopheles coluzzii*) detect and escape from a mechanical swatter (Fig. 1). We did this using two sets of experiments (Figs 1c-e and 2-4, respectively). In the first set, we quantified the relative contribution of attacker-induced visual cues and airflow to the escape performance of night-flying mosquitoes. In the second set, we investigated how these mosquitoes escape from the attacker, and how they use the attacker-induced airflow actively and passively.

**Fig. 1:**
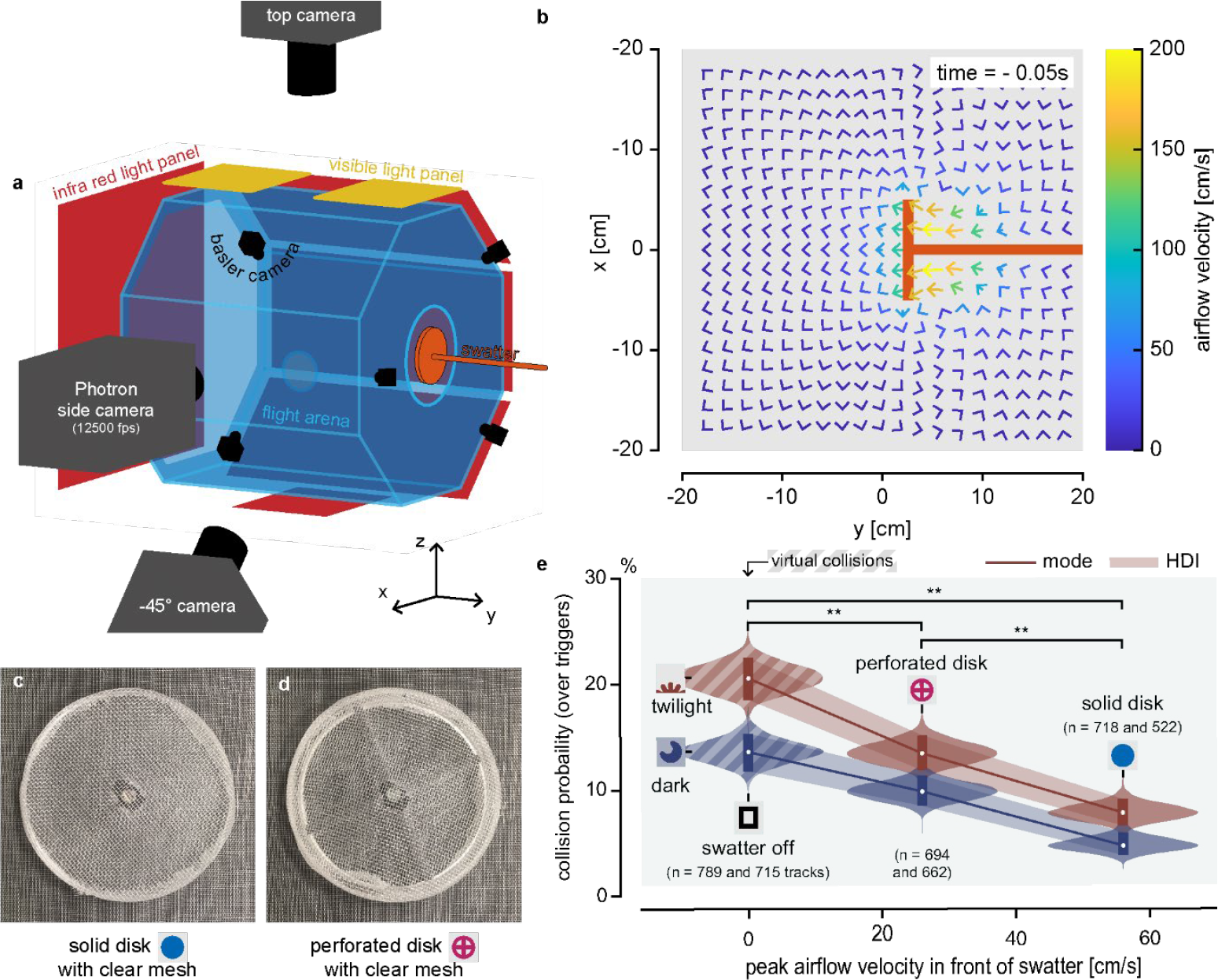
Experimental setup and mosquito collision probability. (**a**) The experimental setup consists of a flight arena, a mechanical swatter and two videography systems (a five Basler camera real-time tracking system and a high-speed videography system with three Photron SA-X2 cameras recording at 12,500 frames per second). All cameras used infrared lights for illumination; visible light panels were set to twilight or turned off. (**b**) Airflow velocity field at time *t*=-5 ms before maximum swatter excitation. (t=0 s) (**c**-**e**) In our first experiment, mosquitoes were attacked by one of two disks covered with a clear mesh: a solid disk swatter (**c**) or a perforated disk (**d**). (**e**) Bayesian estimate of collision probability as function of light condition, and peak airflow induced by the swatter (measured using hotwire). For the swatter turned off, we estimated virtual collisions from tracked positions of mosquitoes and numerically simulated swatter movement. ** show statistically significant differences (see (10) for details); HDI: 89% Highest Density Interval; PDF: Probability Density Function.

## Results and discussion

In our first set of experiments, we investigated whether malaria mosquitoes rely on air movement to escape from a looming threat. To do so, we tracked free flying mosquitoes in real time and – based on their position and velocity – we triggered a mechanical swatter to simulate an attack towards the mosquito’s predicted position (Fig. 1a, S1). To vary the airspeeds generated by the swatter, we used two swatter disks (Figs 1c and 1d): one solid and one perforated, producing high and low swatter-induced airflow, respectively. To test the effect of visual cues on escape performance, we performed these experiments both in the dark and in twilight. A control with the swatter turned off produced no airflow and no visual cues. Using Bayesian statistics, we found that the probability of a collision between swatter and mosquito is lower in the dark (Fig. 1e); a previous study showed that this is due to an increased flight path unpredictability with reducing light intensity (18). Moreover, in both light conditions, collision probability scaled negatively with increasing swatter-induced air movement. This shows that mosquitoes use the swatter-induced air movement to escape the looming threat, both in the dark and in twilight.

Thus, escaping mosquitoes may use air movements as sensory cues to actively trigger their escape, or they may passively move with the airflow to avoid collision. Our second experiment aimed at determining the relative contribution of these active and passive mechanisms to their escape performance. For this, we augmented our real-time mosquito tracking system with a high-speed stereoscopic videography system (Fig. 1a, Movies S1-S3). We then applied a custom-made deep-neural-network tracking algorithm to the recorded videos to track the body and wingbeat kinematics of mosquitoes escaping the swatter in twilight (Fig. 2, Movie S4). In this experiment, we also varied the visual cues that were produced by the swatter by using either an opaque or transparent swatter disk (Fig. 1g). By combining experimental hotwire anemometry measurements with computational fluid dynamic (CFD) simulations (Fig. 1b), we quantified the relative airflow velocity in the mosquito reference frame (Fig. 2f,g). Using the tracked kinematics, the CFD results and a mechanistic movement model of escaping mosquitoes, we estimated how these flying animals make use of active and passive escape mechanisms to evade fast approaching objects in low light conditions (Figs 2-4).

**Fig. 2:**
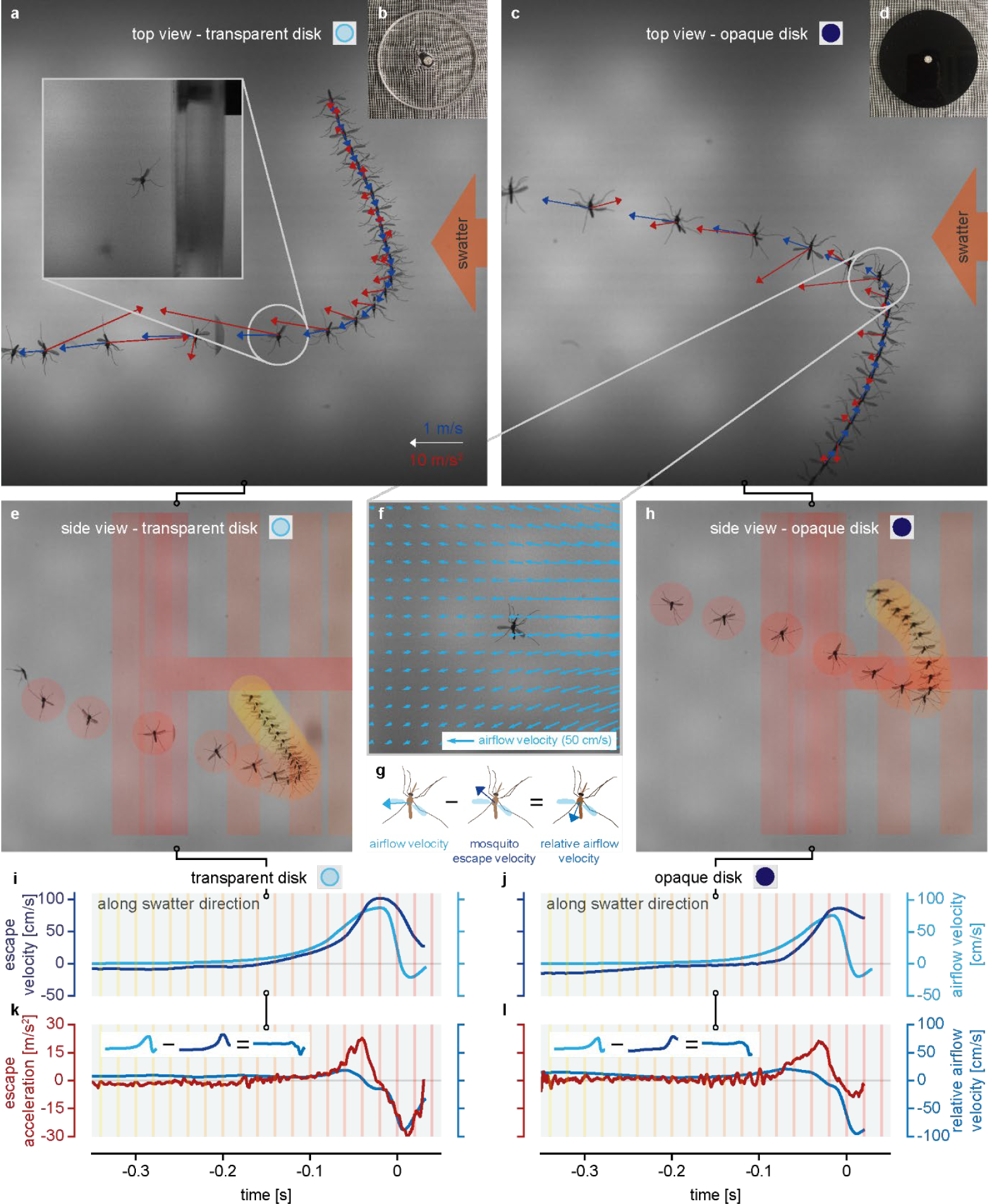
Escape manoeuvres of two mosquitoes attacked by a transparent (b) or an opaque swatter (d). (**a**,**c**,**e**,**f**) Photo montages of the escaping mosquitoes with various data overlayed: (**a**,**c**) instantaneous velocity (blue) and acceleration vectors (red); (**e**,**h**) mosquito and swatter positions color-coded with time (defined in (**i**-**l**)). (**f**) Instantaneous airflow velocity field produced by the swatter around the mosquito (from CFD). (**g**) scheme defining the relative airflow velocity. (**i**,**j**) Temporal dynamic of the escape velocity and swatter-induced airflow velocity at the mosquito’s position. The swatter reached is most forward position at t=0 s. (**k**,**l**) Equivalent escape acceleration and relative airflow velocity in the mosquito reference frame, as defined in (**h**).

Results show that mosquitoes escaped from the looming swatter by first performing a rapid turn manoeuvre away from the approaching object, and then continued to fly away from it at an increased flight speed (Fig. 2a,c, Movies S1-S3). To produce both the rapid turn and the increase in flight speed, the escaping mosquitoes exhibited one or sometimes several high acceleration peaks away from the incoming swatter.

All the mosquitoes that escaped successfully were heading away from the attack at the end of the swatter movement (Figs 3, S14). However, due to differences in initial flight direction, the turn kinematics varied between manoeuvres (Figs 3, S15). Mosquitoes attacked from the back exhibited the smallest escape (ground) velocities and accelerations, whereas mosquitoes attacked from the front exhibited the highest escape velocities and accelerations (Fig. S15*c,e*). On average, mosquitoes had a nearly constant body yaw during the manoeuvres and similarly rolled toward the side opposite of the attack (Fig. 3c,e). In contrast, they exhibited very different body pitch kinematics (Fig. 3d). Mosquitoes attacked from their side exhibited on average very little changes in body pitch, whereas mosquitoes attacked from the front and back pitched up and down, respectively.

**Fig. 3:**
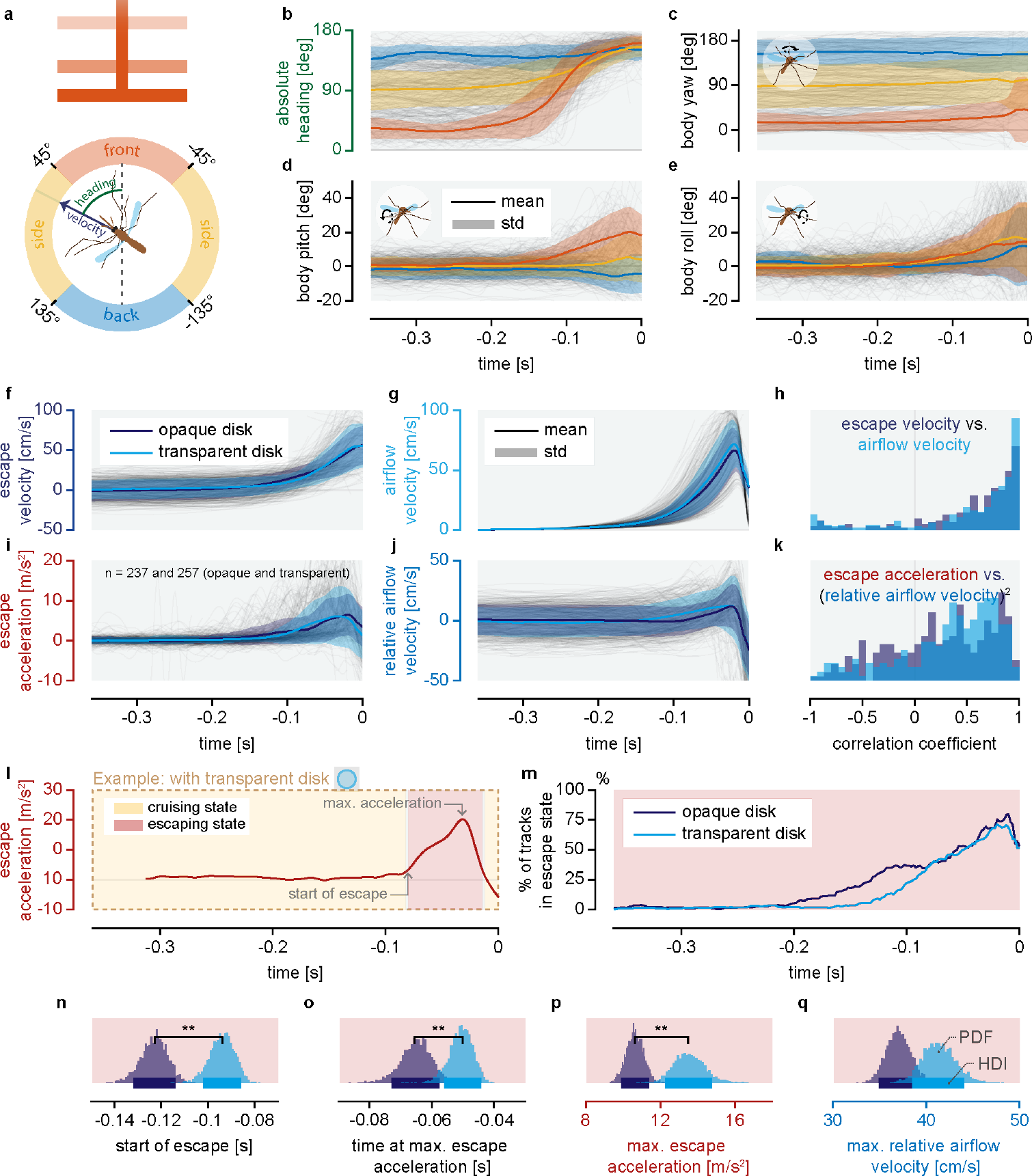
Flight kinematics of mosquitoes escaping from a mechanical swatter. (**a**,**b**) Mosquito initial heading is defined as the mean angle between mosquito velocity vector and the direction of the swatter between -0.32 and -0.24 seconds before the swatter reaches its most forward position. (**c**-**e**) Dynamics of mosquitoes’ body tilt angles in function of their initial heading. Around half of mosquito manoeuvres have been mirrored in order for their roll angle to be positive when they rolled away from the swatter (i.e. as if mosquitoes are all flying from the same side). The body yaw, pitch and roll are defined using Tait-Bryan convention. (**f**,**g**,**i**,**j**) Temporal dynamics of (**f**) escape velocity, (**g**) airflow velocity at the mosquito’s positions, (**i**) escape acceleration, and (**j**) the relative airflow velocity in the mosquito reference frame, for all recorded escape manoeuvres from the opaque swatter (purple) or transparent swatter (blue). Grey lines show individual tracks. (**h**,**k**) Histograms of the correlation coefficients for all tracks: (**h**) between escape velocity and airflow velocity; (**k**) between escape acceleration and relative airflow velocity squared, as an airflow-induced drag scaling parameter. (**l**,**m**) Quick escape manoeuvres were identified using a Hidden Markov Model (HMM): (**l**) Temporal dynamics of the escape acceleration of a flying mosquito, including the cruising or escaping states identified using our HMM. (**m**) Temporal dynamics of the escape state proportions. (**n**-**q**) Bayesian estimates of various escape performance metrics: (**n**) Starting time of the escape state; (**o**) time of maximum escape acceleration; (**p**) maximum escape acceleration; (**q**) maximum relative airflow velocity. ** show statistically significant differences; HDI: 89% Highest Density Interval (see (10) for details); PDF: Probability Density Function.

Previous research showed that flying mosquitoes comply well to the so-called helicopter model for flight control (25). Given this steering mechanism, mosquitoes escape from a looming threat by performing a banked turn. Hereby, the roll kinematics and the consequent sideways force production is independent of the circumferential location of the looming object. In contrast, the pitch kinematics depends strongly on looming object location, whereby pitch rotations cause the animal to produce an aerodynamic thrust force away from the approaching danger. This includes pitch up and down rotations when the danger comes from the back and front, respectively. For cases when the swatter looms from the side, a small pitch up rotation is observed. Finally, yaw kinematics do not seem controlled at all, causing large sideslip at the end of each escape manoeuvre.

Mosquitoes escape kinematics are similar to those observed in fruit flies responding to a visual looming stimulus, with some notable differences (20, 23). Fruit flies vary both the roll and pitch kinematics based on looming location, whereas mosquitoes only vary pitch kinematics. The pitch kinematics also differs between the Diptera species, where fruit flies always pitched up and mosquitoes perform both pitch up and down manoeuvres. Finally, fruit flies partly realign their yaw angles with their headings using a passive aerodynamic coupling mechanism (20, 35), whereas the escaping mosquitoes tended to hardly rotate around yaw axis. This suggests that the passive yaw control mechanism identified in escaping fruit flies (35) is less effective in escaping mosquitoes, and that they would need to actively correct their yaw later when the danger has already been avoided.

The body rotations observed in the escaping mosquitoes might be the result from both passive airflow-induced aerodynamic torques and torques produced actively by the mosquito. The fact that mosquitoes attacked from the front and back produce similar roll kinematics as those attacked from the side suggests that at least these roll rotations are for a large part actively induced (Fig. 3e).

Next, we studied how the body rotations and swatter-induced airflow resulted in variations in velocities and accelerations throughout the escape manoeuvres (Fig 2). We found that the escape velocities align well with the surrounding airflow velocity (Figs 2i,j and 3f-h). This shows that mosquitoes escaping from the swatter tend to travel with the induced airflow. Similar airflow usage has previously been observed for traveling long distances with the wind (36).

If the escapes of mosquitoes were fully passive, the peak accelerations at the start of the manoeuvre would have resulted from the swatter-induced aerodynamic drag on the mosquito. Such a drag force scales with the relative airflow velocity squared, in the mosquito reference frame (*U*_air,rel_^2^) (Fig. 2g). However, the observed escape accelerations correlate poorly with *U*_air,rel_^2^ (Figs 2k,l and 3i-k). This shows that the high acceleration peaks produced during the start of the escape manoeuvres are primarily actively produced. In contrast, the decelerations during the second phase of the escape manoeuvre align well with the relative air speed squared (Fig 2k,l), showing that in this part of the escape manoeuvre the mosquitoes tend to travel passively with the bow-wave produced by the swatter.

Comparing the escapes from the opaque or transparent swatters shows that mosquitoes attacked by the opaque disk started their escape earlier (Fig. 3m,n). Also, they reached their peak accelerations earlier, but with lower peak amplitudes (Fig. 3o,p). This shows that mosquitoes rely partly on visual cues to trigger their escape manoeuvres, as was previously suggested (18, 19). It also confirms that the peak accelerations at the start of the escape are produced actively by the mosquitoes.

We subsequently used our mechanistic model of a mosquito escaping from a swatter to estimate the relative contribution of the active and passive mechanisms to the overall escape performance (Fig. 4a-c, S19). Here, we modelled the aerodynamic forces acting on an escaping mosquito as the sum of its active wingbeat-induced forces and the passive swatter airflow-induced forces (Fig. 4d-f). In our model, the magnitude of wingbeat-induced force depends on the product of the wing stroke amplitude and frequency, and the force direction depends on body orientation. Both the increase of the total aerodynamic force and body tilt angle correlate with increases of the wingbeat amplitudes and frequencies during the manoeuvres (Fig. 4c and Fig. S17*c,d*). These wingbeat kinematic parameters are unlikely to have been changed by the airflow induced by the swatter. This provides further evidence that, during the escapes, mosquitoes are actively increasing the aerodynamic force on their body.

**Fig. 4:**
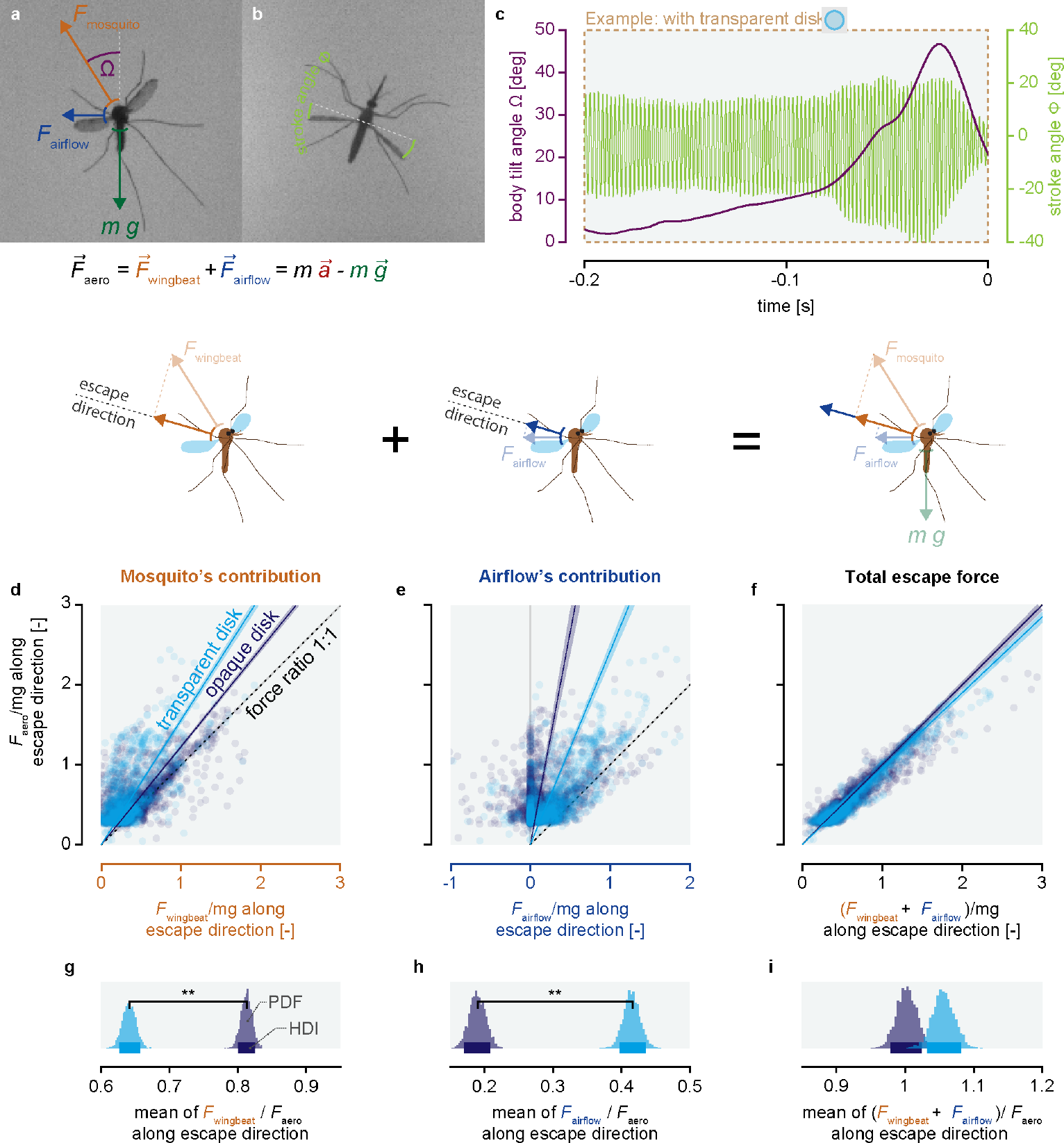
Wingbeat-induced forces and swatter airflow-induced forces acting on escaping mosquitoes. (**a**,**b**) Schematics describing body tilt angle (Ω), wing stroke angle (Φ), and the forces acting on the flying mosquito: body weight *mg*, and the aerodynamic force (*F*_aero_) that consists of the wingbeat-induced force (*F*_wingbeat_) and swatter airflow-induced force (*F*_airflow_). (**c**) Temporal dynamics of body tilt and wing stroke angle of a mosquito escaping from the transparent swatter. (**d**-**i**) The relative contribution of *F*_wingbeat_ and *F*_airflow_ to *F*_aero_ in the escape direction, for escapes from the opaque and transparent swatter (in purple and blue, respectively). All forces are normalized with weight and from mosquitoes in the escaping state. (**d-f**) Total aerodynamic force (ordinate) as a function of (**d**) the wingbeat-induced force, (**e**) swatter airflow-induced force, and (**f**) their sum. Purple and blue lines with bars show Bayesian estimated linear fits. (**g-i**) Bayesian estimates of the contribution of each force component to the total aerodynamic escape force (equal to the slope inverse in (**d**-**f**)). ** show statistically significant differences (see (10) for details); HDI: 89% Highest Density Interval; PDF: Probability Density Function.

By applying our model to all 494 tracked escape manoeuvres, we estimated the relative contribution of the active wingbeat-induced forces and passive swatter airflow-induced forces to the total aerodynamic forces in the escape direction (Fig. 4d-i). We found that when the swatter is relatively well visible (opaque disk), most of the acceleration away from the looming object is actively generated (81%, Fig. 4g). Mosquitoes generates this active acceleration by rotating their bodies away from the danger and by increasing their wingbeat amplitudes and frequencies (Fig. 4c and S17).

Even when almost no visual cues are available (transparent swatter in twilight), mosquitoes still actively produced more than half the aerodynamic forces required for their escape (64%, Fig. 4g). This suggests that nocturnal mosquitoes can detect an attack without using vision, presumably by sensing the airflow induced by the attack. For this, they might rely on aerodynamic imaging (37), although aerodynamic imaging has a range limited to a couple of body lengths, and many of the escaping mosquitoes never got that close to the swatter (Movie S3). Alternatively, mosquitoes may directly detect the minute increases in airflow that precedes the swatter, or detect airflow-induced body rotation using their halteres (34, 38).

Both when mosquitoes had relatively high or almost no visual information about the attack, the passive contribution of the airflow in their escape force was far from negligible (19% and 41%, respectively). Thus, our results show that mosquitoes escaping from a looming object detect the danger using combined visual and non-visual cues. They then actively respond by rapidly accelerating away from the swatter, and finally continue to move away from the danger by travelling with the swatter-induced bow-wave.

Although the passive effects of the air moved during an attack are unlikely to play a significant role in the escapes of big insects, it is not surprising that airflow affects insects like mosquitoes as they have low mass and inertia (25). Among the millions of flying insect species, mosquitoes are of average size (39), and so the majority of flying insects could make use of similar looming object-induced airflow to escape attacks or avoid collisions.

Because passive use of airflow has been suggested as an important driver of the evolution of controlled powered flight in insects (40), studying how insects passively use airflows to escape might also help us understand how their control system evolved. Finally, our results help explain how mosquitoes are able to escape from the inward airflow generated by odour-baited traps (41), and consequently, this provide new insights on how to improve these traps in order to better control mosquito populations.

## Materials and methods

### Experimental animals

For our experiments, we used non-blood-fed adult *Anopheles coluzzii* females (age = 7.4 ± 1.2 days post emergence (mean ± std)). These mosquitoes came from a colony that originated from Suakoko, Liberia in 1987. This colony was reared in a climate chamber inside the Laboratory of Entomology (Wageningen University & Research, The Netherlands) where the temperature and relative humidity were respectively kept at 27°C and at 70%. The light cycle inside the room consisted of a shifted clock with 12h light and 12h dark periods. Adult mosquitoes were reared in BugDorm cages (30 × 30 × 30 cm, MegaView Science Co. Ltd., Taiwan), where they had constant access to 6% glucose sugar water solution. Mosquitoes were divided in two groups, either to be used for experiments or to be used for rearing. Rearing mosquitoes were blood-fed daily with human blood (Sanquin, Nijmegen, The Netherlands) using a membrane feeding system (Hemotek, Discovery Workshop, UK). These mosquitoes had access to wet filter papers upon which females could lay their eggs. After being collected, these eggs were moved into plastic larval trays filled with 27°C water containing a few drops of Liquifry No. 1 fish food (Interpet, UK). After emerging, the larvae were fed with TetraMin Baby (Tetra Ltd, UK). To emerge, pupae were moved into new BugDorm cages. Both males and females were kept together.

### The flight arena

To understand how mosquitoes escape from being swatted, we filmed them while freely flying in an octagonal flight arena (50×50×48cm (height x width x length)) with transparent Plexiglas walls (Fig. 1A)(18). In the front and back of the flight arena, there were two circular holes closed by HFPE insect screening (Howitec, The Netherlands) to allow air circulation and facilitate cleaning (18). The temperature and relative humidity inside the arena were controlled by the climate control system of the room (42). To provide visual cues for mosquitoes, visual markers (randomly shaded grey squares printed on paper) were placed on the floor of the flight arena. Also inside, a sensor (AM2302, ASAIR) was recording relative humidity and temperature.

A visible light LED panel was positioned above the arena to change the light condition from dark (turned off) to twilight or sunrise (Fig. 1A and S3). To mimic twilight condition, multiple polyester neutral density filters of 0.8 ND (LEE filters, Panavision Inc.) were placed in front of the LEDs in order to reduce light intensity. Additionally, the flight arena was surrounded by multiple infrared LED panels. Because mosquitoes cannot see infrared light (43), we could track them in the dark using five infrared enhanced cameras (Basler acA2040-90umNIR) with 12.5 mm lenses (Kowa LM12HC F1.4) and the real-time tracker Flydra (version 0.20.30) (44, 45). These cameras filmed mosquitoes at 90 frames per second and at a resolution of 680×680px (with a pixel-binning of 3). Lens distortions were corrected by filming a backlighted print of a chequerboard pattern.

We simulated attacks using a mechanical swatter made of a 1 cm diameter black aluminium shaft and an opaque or transparent plexiglass disk. This disk had a size similar to a human hand with a diameter of 10 cm and a thickness of 1 cm. The swatter was moved along a 50 cm long toothed belt axes (drylin ZLW-1660-G0BW0-D0A3B-0A0A-500) powered by an AC servo motor (Schneider Electric Lexium BCH2 LD0433CA5C), itself controlled by a programmable motion servo driver (Schneider Electric Lexium LXM28A). The swatter kinematics, similar to a human swatting or a bat attack(30), was the same as in our previous study(18). The swatter was triggered based on real-time prediction of mosquito position 367.5 ms in the future (i.e. when the swatter would be around halfway to its final position, t = -157.5 ms in Fig. 1-4). These predictions were linear estimations based on mosquito’s current three-dimensional position and velocity (assumed constant for the prediction). Thus, the swatter was triggered if a mosquito was predicted to be in a 10 cm diameter spherical triggering region at the centre of the flight arena. After being triggered, and having waited one second, the swatter was moved slowly back to its initial position. After a delay of ten seconds another trigger was allowed. Finally, during post-processing, we filtered out all the triggers for which the mosquitoes were not predicted to be inside the sphere of interest when the swatter would reach its most forward position (t = 0 s).

Two different experiments are presented in this study, the first was carried out to investigate the effect of air gusts on the escape performance of mosquitoes (Fig. S1). For this, we used two types of transparent disks, a solid and a perforated disk, which generated different amounts of air movement (Fig. S2). In order to control the effect of visual cues, a clear mesh (Ornata plus 95135, howitec.nl) covered both sides.

The main goal of the second experiment was to zoom in on mosquito escapes in order to record in detail their body and wing kinematics during those manoeuvres. For this, three high-speed cameras (FASTCAM SA5, Photron) were added to the setup. These cameras were pointed toward the centre of the flight arena and recorded images at 12500 frames per second and at a resolution of 1024 × 1024 pixels. They filmed a 12×12×12 cm volume around the final position of the swatter, where mosquitoes were initially predicted to fly towards. Several small modifications of the setup were made to allow the addition of the high-speed cameras (Fig. 1A). These consisted of the removal of a part of the visible light panel above the arena, the addition of an infrared light panel below the arena, and the decentring of a hole in the plexiglass side panel to plug in a mosquito release cage. Finally, to reduce the chance of having individual mosquito out of the focus plans of the high-speed cameras, the triggering parameters were slightly modified. These involved reducing the size of the triggering region to a 5 cm diameter sphere and in lowering the latency used to compute the predicted positions to 315 ms (*t* = -0.210 s on Fig. 2-4).

### Experimental procedure

The day before each experimental night, a disk was attached to the swatter rod according to a quasi-randomized planning (Fig. S4*a*). The experimenter cleaned the flight arena with paper towels and a 15% ethanol solution. Skin odour contamination was avoided by handling all materials and mosquitoes while wearing nitrile gloves. Then, a calibration of the Basler cameras was done by tracking the position of a manually waved single LED in the arena. The calibration was aligned using an alignment device with eight LEDs at known three-dimensional coordinates. Calibration snapshots were taken with the Photron cameras of a calibration device with 23 3.5 mm beads at various known three-dimensional positions. These snapshots were later used to compute DLT coefficients for each camera. After closing the flight arena, a release cage with 50 female mosquitoes was plugged to its side and mosquitoes were released to fly freely inside. The experimental procedure was then automatically controlled by a Python 2.7 script. This script allowed for the automatic changes of the light condition inside the flight arena, the tracking of mosquitoes and the triggering of the swatter. For the first experiment, trials started the following morning at 2:30 a.m., two hours after the start of mosquitoes normal dark phase. Three different light conditions were tested consecutively per night, each for 160 minutes. The order of these light conditions was changed following the previously mentioned quasi-randomized planning (Fig. S4*a*). For the second experiment, only the twilight condition was tested, and the experiment lasted through the entire dark phase of mosquitoes (0:30 a.m. to 0:30 p.m.) (Fig. S4*b*). In the afternoon, mosquitoes were removed from the arena using a vacuum cleaner, and left inside its bag to desiccate.

### Analysis of three-dimensional flight tracks

We did the pre-processing of the first experiment dataset using Python 2.7. First, collisions were identified by looking at the two-dimensional tracking results of the Basler cameras positioned at the side of the flight arena. Then, the two-dimensional points corresponding to the swatter were filtered, and three-dimensional tracks were reconstructed again. To be able to compare the chance of being hit by the swatter with or without a disk, collisions with a virtual swatter were determined. These virtual collisions were estimated by computing if and when mosquito flight tracks would have crossed the path of a virtual disk. Then the data was analysed using Matlab R2019b. Outliers of computed three-dimensional points were filtered out using the covariance matrices estimated by the Flydra tracker extended Kalman filter. Segments that were shorter than 4 points long (at 90 fps) were filtered out and segments that were separated by more than 15 points were divided into separated tracks. Then, missing values were interpolated (makima, Matlab) and smoothed using a Savitzky-Golay filter with a moving window of five frames. Finally, for the rest of the analysis, we only kept the tracks that started at least 60 frames before and 30 frames after the time at which the swatter reached its most forward position (t = 0s on all the figures).

The second dataset (using the high-speed cameras as shown in Fig. 1) was also analysed using Matlab R2019b. Only the tracks that did not end up in a collision (if mosquitoes touched the disk with any body part) were kept for further analysis. Three-dimensional track segments that were shorter than 50 frames (at 12500 fps) were filtered out. Similarly, segments that were separated by more than 20 frames from the main track were also filtered out. Instantaneous wingbeat amplitudes and frequencies were estimated using a Hilbert transform on the wings’ stroke angles. Then body and wings parameters were smoothed using a Savitzky-Golay filter with a moving window of 251 frames. After that, framerate was lowered to 500 fps (linear resampling) for the rest of the analysis.

For both datasets, flight velocities and accelerations over time were computed using central finite difference schemes (with an accuracy of 6). Initial and final values were estimated respectively using forward and backward finite difference schemes. Then, the escape velocity and escape acceleration were defined and computed by projecting mosquito velocity or acceleration over time on the moving line between the nearest point on the swatter and mosquito three-dimensional position (see Supplementary Fig. 12*c*).

### Quantifying body and wing motion

To investigate mosquitoes’ reaction to the swatter attack, it was necessary to accurately estimate their position as well as their body and wing angles throughout the manoeuvres. Tracking the body parts of mosquitoes from our dataset was challenging. Indeed, due to the relatively large three-dimensional volume filmed and to experimental lighting limitations, body resolution was low and contrast was restricted. This meant that classical automatic tracking methods would most likely yield poor results. Manually tracking body parts was also not an option because our dataset comprised of several hundreds of thousands of images. Thus tracking needed to be automated with an accuracy close to human tracking. To accomplish this goal, we developed a new flying insect tracker written in Python 3.6 and based on the deep neural network package DeepLabCut (46). This open-source package uses transfer learning to do markerless pose estimation using relatively low numbers of labelled images.

The tracking process is described in detail in the supplementary Fig. S5. It starts with preparing low-resolution videos to be used by DeepLabCut. First, mosquitoes are tracked in two-dimensions on each of the three different camera views using the opencv2 blob detection algorithm. Then, to filter out noise, three-dimensional tracks are reconstructed using previously computed DLT coefficients and the tracks are re-projected in each camera two-dimensional views. From these two-dimensional coordinates, the original images are cropped around mosquitoes’ positions over time. These cropped images are stitched together and converted into a single .avi video per manoeuvre. Such video presents several advantages when compared to the original recordings to be used by DeepLabCut, namely of having a greatly reduced resolution, thus increasing the tracking speed. And it gathered all camera views together, thus allowing the network to learn positional correlation between the views. Using the DeepLabCut GUI, we trained a deep neural network based on the pre-trained network resnet50 and 230 manually labelled images of escaping mosquitoes. For each camera view, we labelled 40 points over the mosquito body and wings, thus resulting in a total of 120 points per image (Supplementary Fig. S6). The deep neural network was then used to automatically estimate positions of these points on all recorded images.

Next, from the two-dimensional results of DeepLabCut we estimated body parameters (i.e. position, lengths and angles of body parts and wings) over time using a custom-made Python 3.6 package. The first steps were to remove outliers with likelihood (provided by DeepLabCut) lower than 0.85 and to reconstruct three-dimensional coordinates of each body part. Then, initial estimation of body parameters was obtained using simple calculus. Because initial estimation of body and wings angles were found to be noisy, updated estimation of these parameters were then obtained by minimizing the sum of squares (scipy.optimize.leastsq) of the root-mean-square deviation between the coordinate of the skeletons joints and the three-dimensional points. Estimated body or wing parameters were filtered out if the root-mean-square deviation was superior to 0.1 mm and if the number of points per wing or body used to do the 3d fitting was lower than 4. Finally, body parameters (but not the wing parameters) were smoothed over time with a low pass Butterworth filter using a cut-off frequency of 100 Hz.

### Classifying tracks as cruising or escaping

To classify flight tracks as either “cruising” (i.e. missing the swatter by chance) or “escaping”, we used a Hidden Markov model (HMM) (Matlab toolbox by Kevin Murphy (47)). In such a model, the studied system is assumed to be a Markov process with hidden states (here cruising or escaping), where the current probability of the system to be in a particular state is only dependent on the previous state and on the chosen output variable(s). Our HMM was trained on all escape accelerations over time of all tracks that didn’t end up in a collision. We estimated the initial parameter of the model by fitting a mixture of two Gaussians (fitgmdist, Matlab) to the distribution of all instantaneous escape accelerations (see supplementary Fig. S7*c*). Then the remaining unknown parameters of the HMM were estimated using a Baum–Welch algorithm, with fixed means and standard deviations (see supplementary Fig. S7*a,b*). The state with the highest mean acceleration was labelled as the “escaping” state while the second was labelled as the “cruising” state. We used the Viterbi algorithm to compute the most-likely corresponding sequence of states for each track (see examples on Fig. 3G). Then we computed the overall proportion of tracks that were in either of the two states for each point in time (Fig. 3H). Finally, we labelled all the tracks as escaping if they were found to be at least once in the “escaping” state (270 tracks). The remaining tracks were labelled as cruising (224 tracks).

### Simulating the swatter-induced airflow

To investigate the role of the airflow generated by the swatter, we simulated this airflow using Computational Fluid Dynamics (CFD) simulation. The simulation of the swatter was performed with the aid of the dynamic mesh solver available in OpenFOAM (version 19.12 (48)), namely overPimpleDyMFoam. The simulations were configured to run with a second order accurate backwards time scheme. The domain consisted of a cylinder (radius = 250 mm and length = 580 mm) with a disk moving in it. An overset (chimera) mesh was employed to enable the movement of the disk, with predetermined kinematics, inside the cylindrical computational domain. This approach combined two distinct unstructured cartesian grids constructed with the aid of CFMesh. A grid was constructed around the disk, and another filled the cylinder domain acting as background grid. Both were predominantly composed of hexahedral cells of 2 mm and 4 mm respectively. No-slip and inletOutlet velocity boundary conditions were imposed at the disk and outer boundaries respectively. Additionally, the simulation used a U-RANS model to account for turbulence effect, more specifically has accomplished that with the aid of a Spalart-Allmaras turbulence model.

The CFD results were validated by comparing them with experiment measurements obtained with a hotwire anemometer (tetso 405i) (see Fig. S8). Two sets of parametric studies were conducted varying the cell sizes and the Courant–Friedrichs–Lewy (CFL) number (Fig. S8 and Fig. S9). The cell sizes of the meshes were selected after comparing multiple mesh sizes to be at the maximum 4 mm for the background mesh and 2 mm for the overset mesh 2 mm. Similarly, various Courant– Friedrichs–Lewy (CFL) number were compared and we selected a final CFL number of 0.4. Because of the chaotic nature of airflow turbulence generated during the movement of the swatter, we could not predict the exact conditions that each individual mosquito experienced. Thus, all of the CFD results presented in this study have been averaged around the axis of the swatter movement (i.e. the three-dimensional axis of symmetry of the swatter) into two-dimensional planes for each millisecond (Fig. S10). Airflow velocities at each instantaneous mosquito three-dimensional positions have been interpolated from these two-dimensional planes using modified Akima piecewise cubic Hermite interpolation (makima, Matlab).

### Estimating active and passive aerodynamic forces during the escape

The escape movement of the mosquito can consist of active and passive components. To estimate their relative contributions to the escape, we modelled their effects on the aerodynamic forces produced during the escape (see free body diagram in Fig. 4A). Here, Newton’s second law of motion dictates that the sum of all force vectors 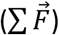 equals body mass times acceleration as

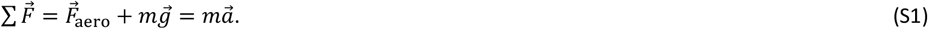

Here, *m* and 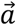 are the mass and acceleration vector of the mosquito, respectively. 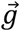 is the gravitational acceleration, and 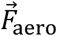 is the aerodynamic force vector acting on the mosquito. This aerodynamic force consists of an active force component generated by the mosquito with its beating wings, and a passive component resulting from the swatter-induced airflow acting on the mosquito. Using a linear approximation, it can be defined as

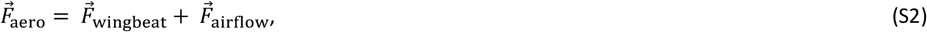

where 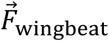 is the active wing-beat induced force vector, and 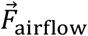 is the passive airflow-induced force vector.

Using a quasi-steady aerodynamic modelling approach (49), we assume that both aerodynamic force components scale linearly with the dynamic pressure and the relevant surface area (*S*) as

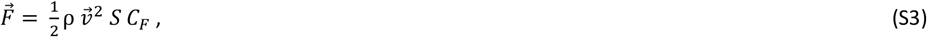

where the dynamic pressure 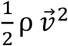 depends on air density ρ and air velocity 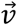. *C*_*F*_ is an unknown force coefficient. We applied this model to both the wingbeat-induced forces and airflow-induced forces.

The wingbeat-induced forces 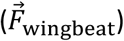 scale quadratically with the velocity of the beating wings (50), and so we estimate the wingbeat-induced dynamic pressure as

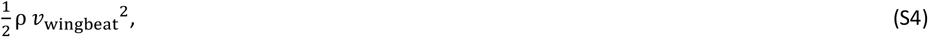

where *v*_wingbeat_ is the average wingbeat-induced speed. By combining equations (S3) and (S4), we can model the wingbeat-induced forces as

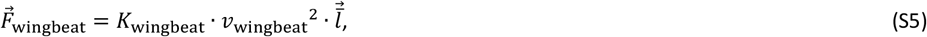

where 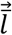 is the unit aerodynamic force vector, and *K*_wingbeat_ is the wingbeat-induced force coefficient that includes all other parameters.

In order to steer, insects following the helicopter model will rotate their body using minute adjustment of their wingbeat kinematics. In our case, this assumption is supported by the fact that no large differences between the kinematics of the left and right wings of mosquitoes can be seen thorough the manoeuvres (Fig. S16). To estimate the direction of the unit aerodynamic force vector, we therefore assume it to remain constant in the body reference frame.

Malaria mosquitoes beat their wings back and forth using a sinusoidal wing stroke motion (25), and thus we can estimate the average wingbeat velocity as

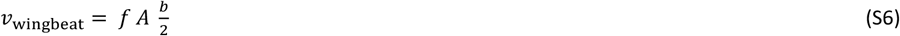

where *f* is the wingbeat frequency, *A* is the wing stroke amplitude, and *b* is the tip-to-tip wingspan (50).

The swatter-induced airflow forces 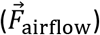 acting on the mosquito scale quadratically with the airflow velocity in the body reference frame of the mosquito. This relative airflow velocity 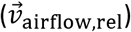 equals the difference between the swatter-induced airflow velocity and that of the moving mosquito as

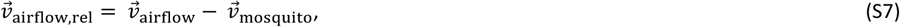

where *v*_airflow_ and 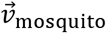 are the velocity of the air and mosquito, respectively. Both 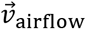 and 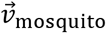 are in the world reference frame, whereas the relative airflow velocity 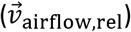 is defined in the mosquito-based reference frame. Based on this, we can now model the airflow-induced aerodynamic forces on the mosquito as

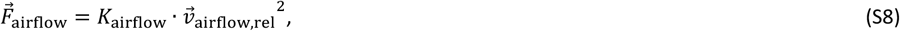

where *K*_airflow_ is the airflow-induced force coefficient that includes all parameters that remain constant throughout a manoeuvre. In this way, *K*_airflow_ would be the same for all directions for an instant in time. This is equivalent to assume that the mosquito body can be approximated by a sphere, and thus that the drag force magnitude on this body will be independent of the wind orientation. Although such assumption may have resulted in estimation errors, we expect these errors to have been maximized along the axis (x and z) where the relative airflow velocities were the smallest, and thus not along the escape direction (along the y axis). Therefore such errors should not have impacted our conclusions.

By inserting the wingbeat-induced and airflow-induced force model equations (S5 and S8, respectively) into our second law of motion equation (S1), and normalizing it with body weight (*m*· *g*), we get

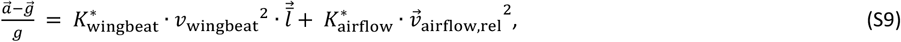

which includes two unknown parameters, being the weight-normalized force coefficients

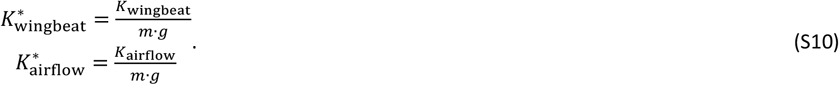

We estimated the magnitude of these two force coefficients for each point in time (i.e. at each digitized video frame) by solving the equation (S9) along the three principal axes using a linear least square solver (lsqlin, Matlab) and constraining the coefficients to be positive. Then, we filtered out estimations with normed residuals larger than 5% of the normalized weight 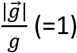.

After estimating these force coefficients, we used equations (S5) and (S8) to compute the temporal dynamics of 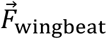 and 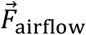 throughout all escape flight maneuvers, i.e. when the mosquito was in the “escape” state. For this, we determined in each escape-state video frame the relative airflow-induced velocity (Eq. S7), and based on the estimated wingbeat frequencies and amplitudes we determined the wingbeat-induced velocity (Eq. S6).

To test the accuracy of our model we first compared the combined airflow-induced and wingbeat-induced aerodynamic forces with the normalized aerodynamic force-induced acceleration 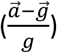 (Fig. 4, S18 and S19). As expected, the sum of airflow and mosquito contributions was close to 100% for both swatters and no significant difference was found between them (Fig. 4F,I). And secondly, because we expected 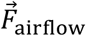to be mostly directed in the horizontal plan, we checked that vertical aerodynamic forces were fully explained by the contribution of 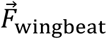 (Fig. S19*g*).

### Statistical analysis

In this study, instead of usual frequentist statistics, we used Bayesian statistics because we think it is conceptually clearer and because it offers richer results by providing estimations of the full-probability distribution of the parameters of interest. We estimated means of mosquitoes’ probability of being hit (Fig. 1E) for each combination of light condition and swatter type using MATJAGS, a Matlab interface for JAGS (51), and the Matlab Toolbox for Bayesian Estimation (MBE)(52), a Matlab implementation of Kruschke’s R code (53). To model the probability of being hit we used a Bernoulli distribution and a logistic link function. Additionally, to compare the escape performance of mosquitoes when attacked by the opaque or the transparent disk, we estimated means of various performance metrics (Fig. 3I-L) and means of the proportion of forces applied to the body of mosquitoes (Fig. 4) using Bayesian estimation (53). We used a normal distribution to model the performance metrics and student t distributions to model the force proportions. Because we had no prior knowledge of the various distributions, we used diffuse priors for the standardized parameters (with a standard deviation equal to 100 times the std of the parameter of interest. Then, we defined standardized effect sizes of the comparison between the two disk types (opaque and transparent) as the difference of their estimated means divided by the norm of their estimated standard deviations.

Finally, we tested for the null hypothesis using the “HDI+ROPE decision rule” (54). For this, we first define the 89% HDI (Highest Density Interval) as the 89% interval in which all the points have a higher probability density than points outside (Fig. S11. And the ROPE (Region of Practical Equivalence) is defined as the range around zero (i.e. the null hypothesis) where a parameter would be found to have “practically no effect”. Therefore, the “HDI+ROPE decision rule” says that the null hypothesis is rejected if the 89% HDI of the standardized parameter (e.g. slopes) completely fall outside the ROPE =[-0.1, 0.1].

## Acknowledgments

We thank Remco Pieters and Henk Schipper for their help in setting up the experiments; Toshi Nakata for helping us in validating our airflow simulation; Guido de Croon, Johan van Leeuwen and Jeroen Spitzen for their feedback during the writing of the article; Cees Voesenek for his advice on animal tracking, and Pieter Rouweler; Kimmy Reijngoudt, André Gidding and Frans van Aggelen for rearing the mosquitoes.

## Author contributions

A.C. and F.T.M. came up with the study. A.C. designed the experiments, build the setup, and performed the experiments. A.C. and P.G. did the airflow measurements. L.H.C. performed the CFD simulations. A.C. analysed the data. F.T.M. provided support for the analysis. A.C. wrote the first draft of the manuscript. All authors contributed to writing the manuscript, and approved the final manuscript.

## Data and materials availability

Original code and data are available in the DRYAD repository https://doi.org/10.5061/dryad.nvx0k6dvz

## Supplementary Materials

Figs. S1 to S19

Movies S1 to S4

Data S1 to S3

Code S1 to S2

