## Supplementary Materials and methods for "Mosquitoes escape looming threats by actively flying with the bow-wave induced by the attacker"

**Consisting of:**

Figs. S1 to S19
Captions for Movies S1 to S4
Captions for Data S1 to S3
Captions for Codes S1 to S2

Movies S1 to S4
Data S1 to S3
Codes S1 to S2

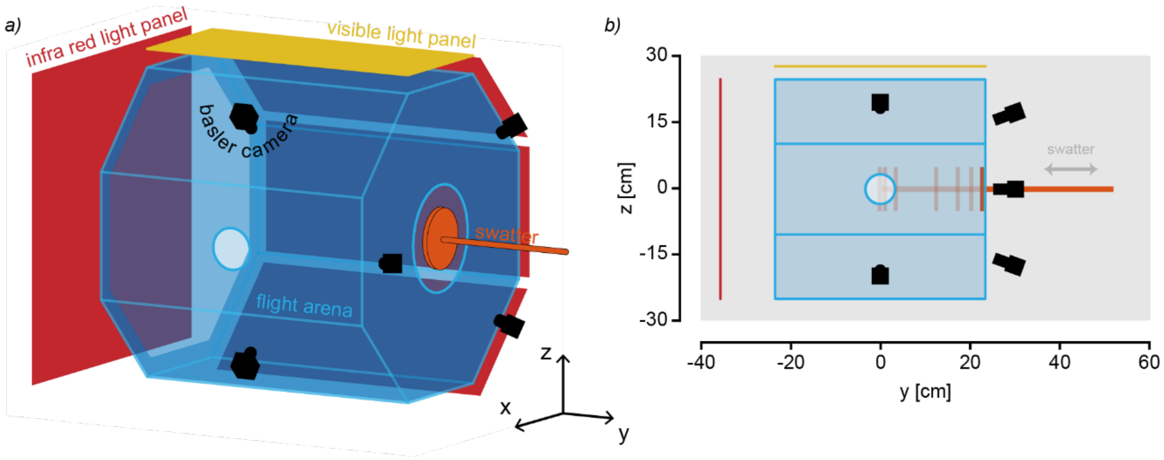


**Fig. S1. Experimental setup to record mosquito flight kinematics.**
(*a-b*) The experimental setup used for the first experiment. In the second experiment, the light panels were modified and the hole to plug a release cage was moved toward the swatter side (see Fig. 1).

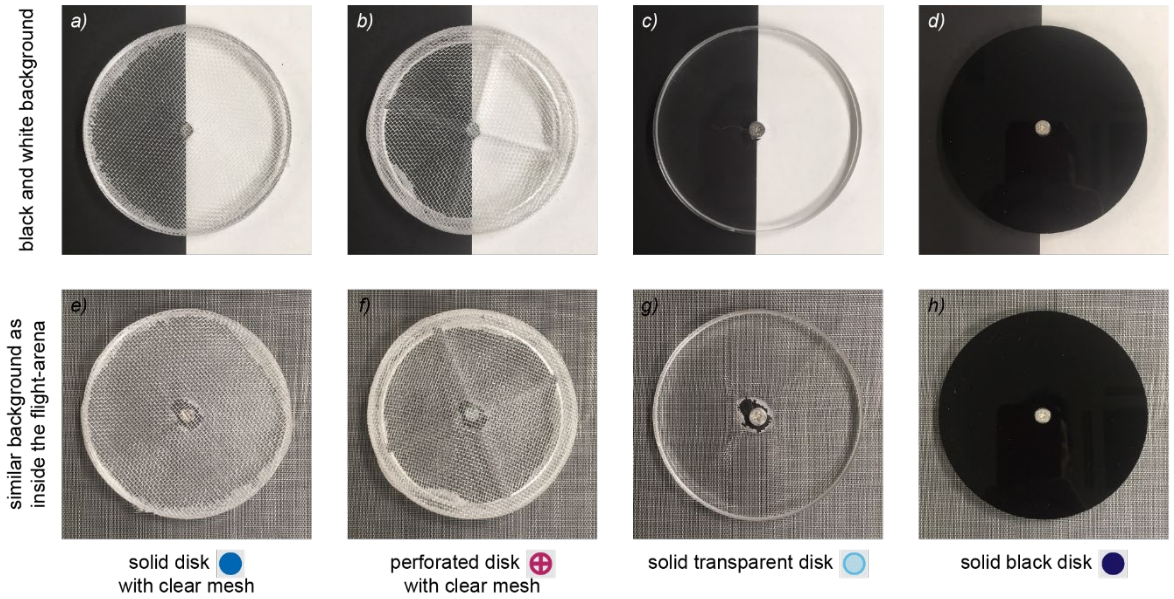


**Fig. S2. Various disks used in the experiments.**
(*a,b,e,f*) Pictures of the solid and perforated disks with a clear mesh that were used in the experiment #2. (*c,d,g,h*) Picture of the transparent and opaque disks that were used in the experiment presented in the Fig. 2-4.

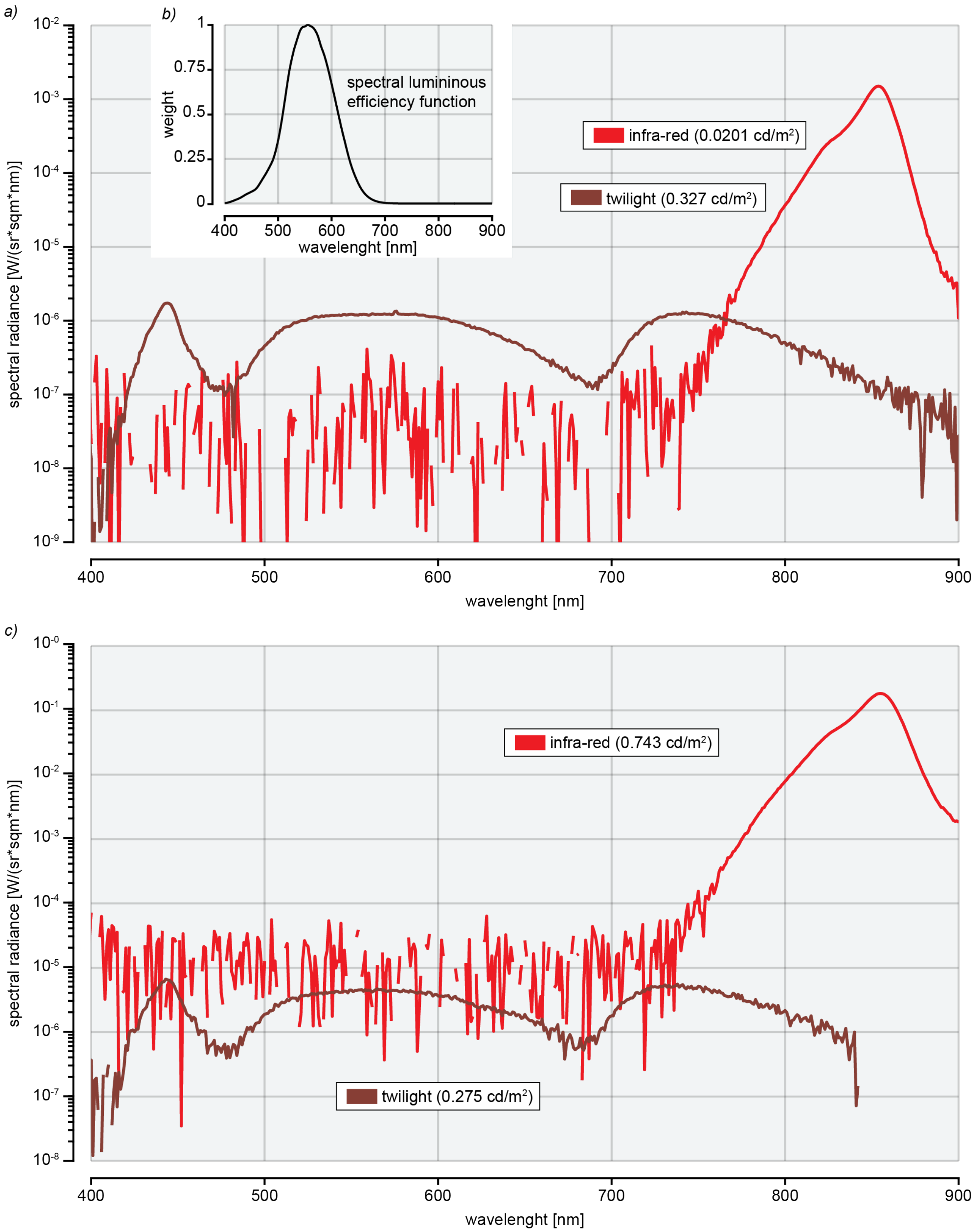


**Fig. S3. Spectrum of the various light conditions.** (*a*) Spectral radiance in log scale of the twilight condition and of the infrared light measured in the middle of the flight arena (as presented in Fig. S2) with a spectrometer (specbos 1211, JETI) and using a diffuse reflector (USRS-99-010-EPV, Labsphere). The low radiance parts of the spectrums appear noisy due to limitations in the range of the radiance that can be measured at the same time. (*b*) Spectral luminous function of humans [1] used to compute the luminance of each light condition. (*c*) Spectral radiance of the light conditions in the experimental setup presented in the supplementary Fig. 1.

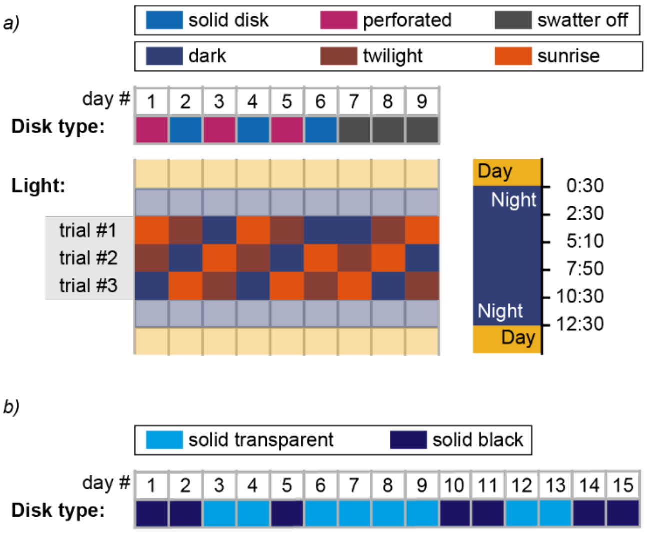


**Fig. S4. Experimental conditions.** (*a*) Experimental planning and conditions for the experiment #1 (results presented in Fig. 1, setup Fig. S2). (*b*) Experimental planning and conditions for the experiment #2 (results presented in Fig. 2-4).

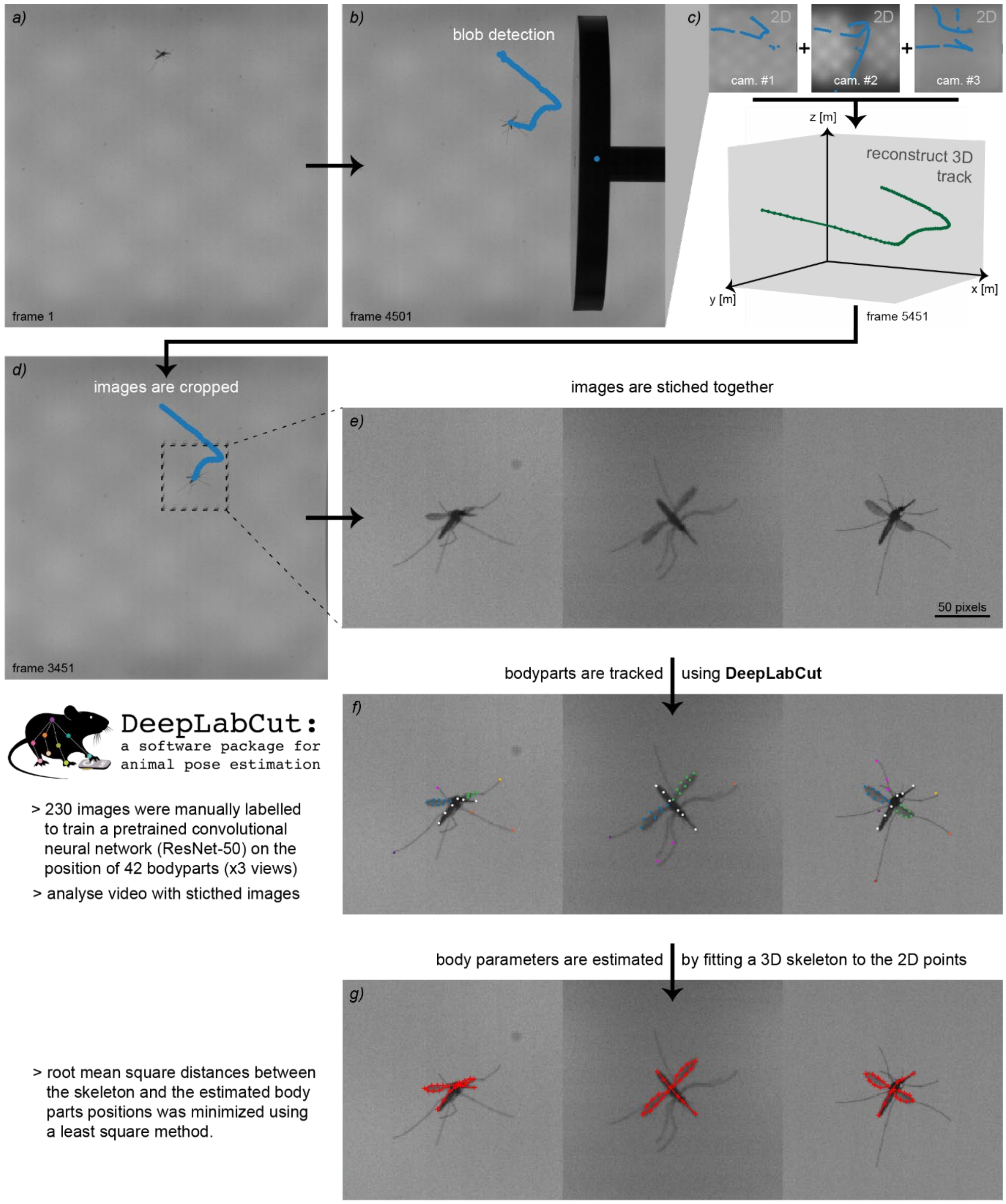


**Fig. S5. Mosquito tracking process.** (*a*) An example of a frame recorded by the side camera while a mosquito was attacked by the mechanical swatter. (*b*) Blobs are detected for all frames of the recording using background subtraction. (*c*) The 2d coordinates resulting from the blob detection on all three camera views are used to reconstruct the 3d tracks of the mosquito. (*d*) The 3d coordinates are then re-projected into each camera view, and the images are cropped around the mosquito positions using a 200 x 200 pixels window. (*e*) The cropped images of the three views are stitched together and converted into a single .avi video. (*f*) The video is analysed using a deep learning network trained with 230 manually labelled images [2]. (*g*) For each frame, a 3d skeleton of a mosquito with variable parameters (e.g. body lengths, span, body and wings angles) is fitted to the reconstructed 3d coordinated of mosquitoes various body parts.

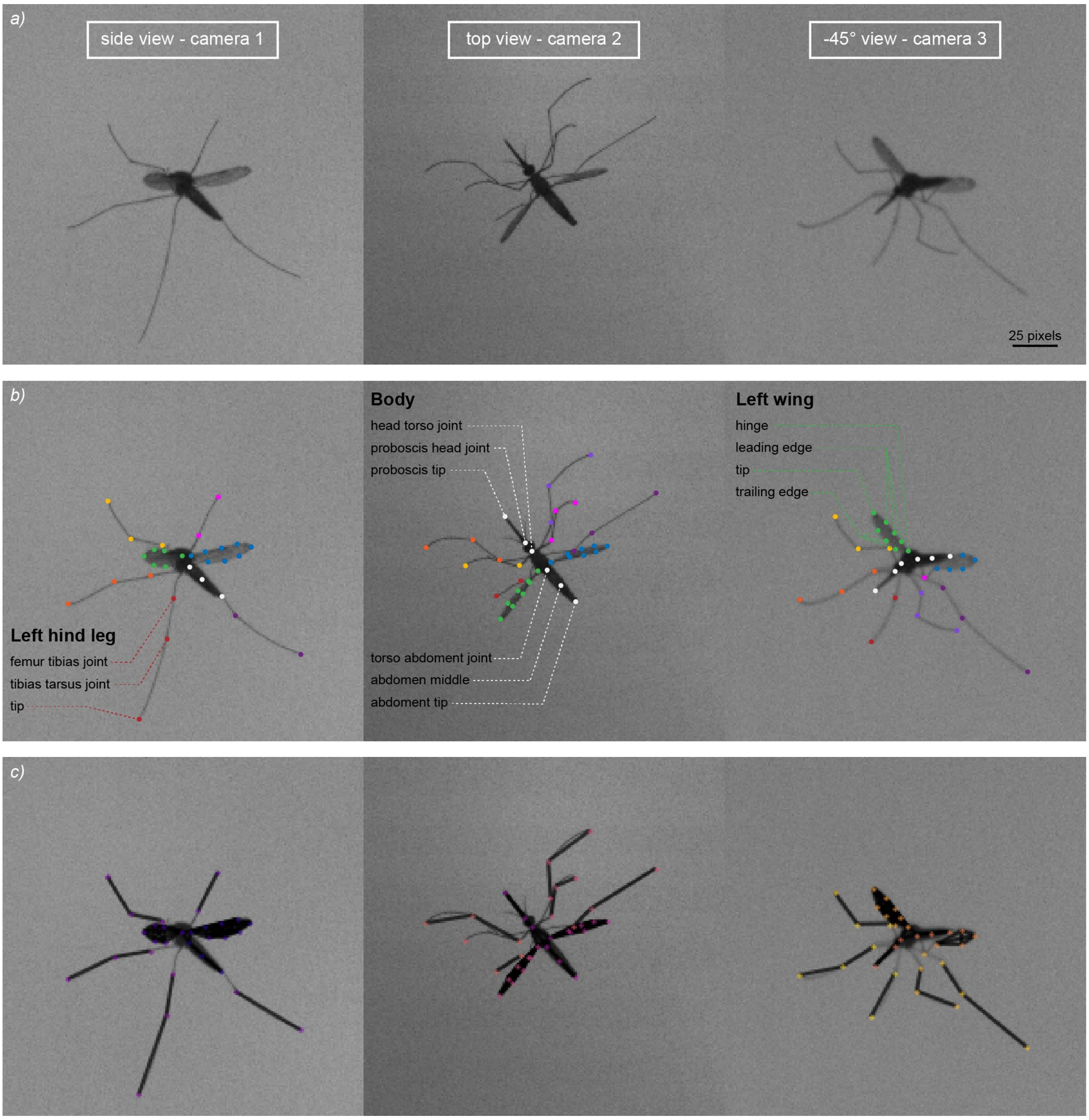


**Fig. S6. DeepLabCut: Body part labels and skeleton.** (*a*) An example of a video frame used in DeepLabCut for the pose estimation of body parts. (*b*) 42 body parts are labelled on each view (x3). (*c*) A skeleton linking the various body parts is defined in DeepLabCut to improve tracking accuracy.

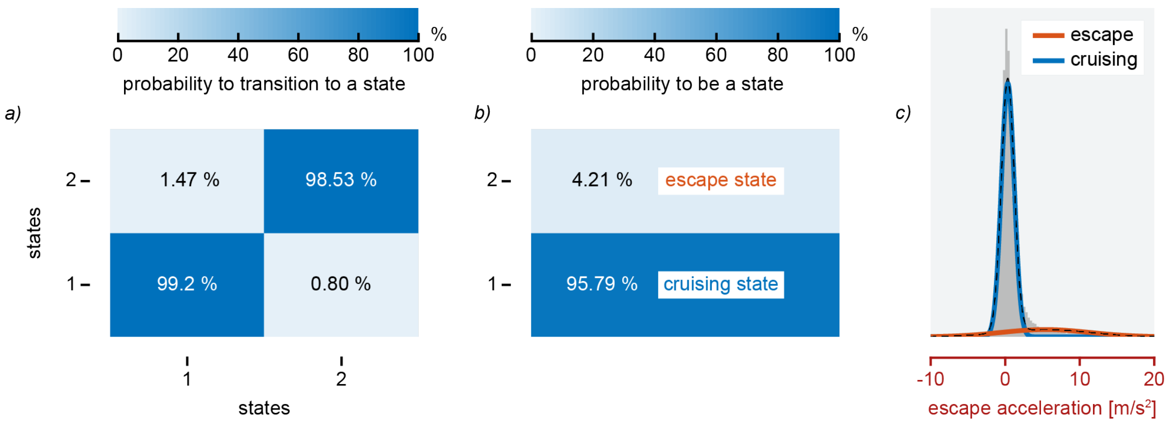


**Fig. S7. Hidden Markov Model parameters.** (*a*) Transition matrix. Each number is the probability of transition from the current state (on the abscise) to a new state (on the ordinate). (*b*) Prior probability to be on each one of the two states. (*c*) ) Distribution of all escape accelerations of the mosquitoes with the Gaussian distribution of the two states used in the HMM.

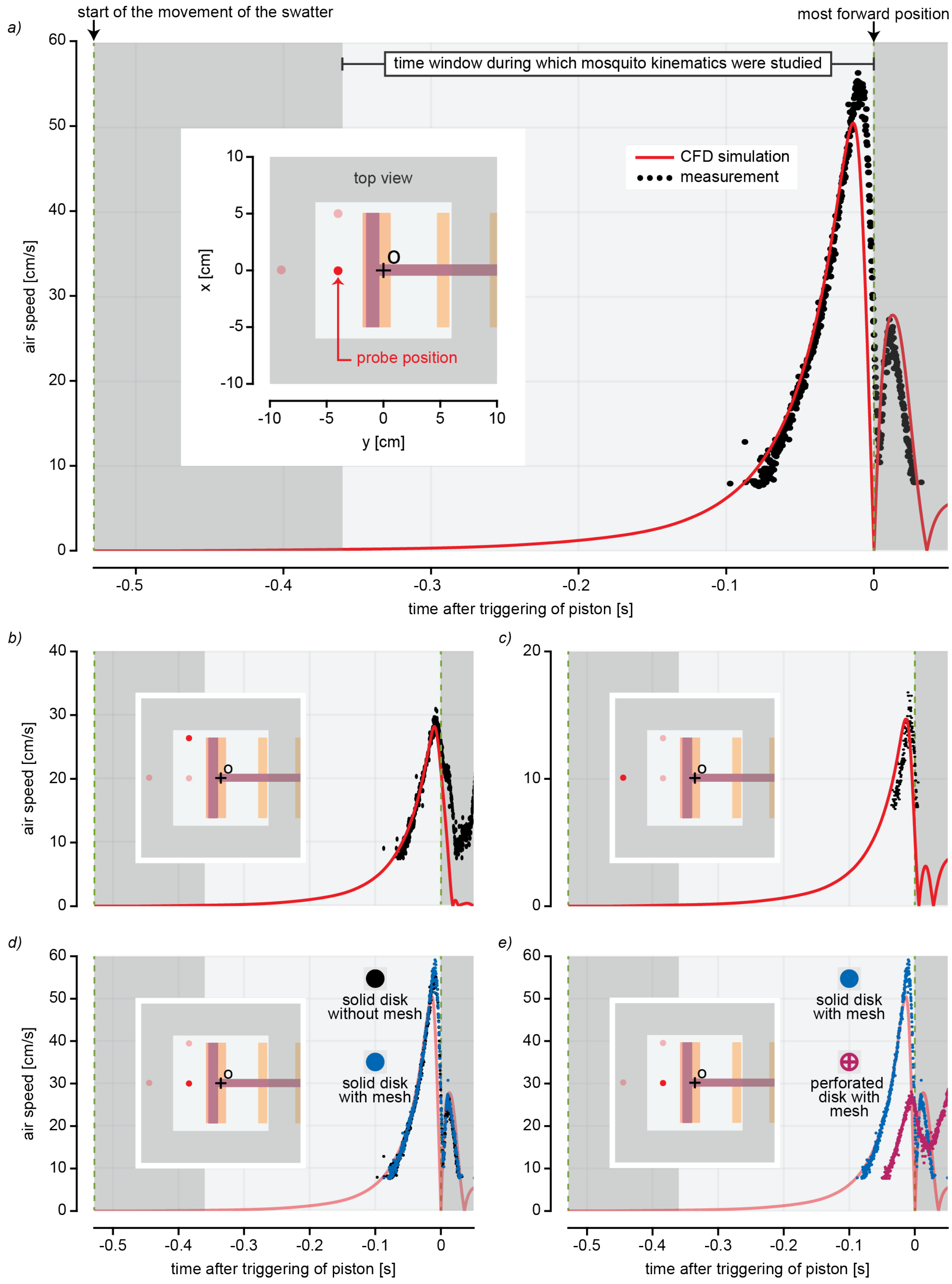


**Fig. S8. CFD simulation validation.** (*a-c*) Comparison between the Computer Fluid Dynamic (CFD) simulation results and the hotwire measurements at three different probe position in from of the swatter with the full disk with mesh. (*d*,*e*) Comparison between hotwire measurements with the three different disk type: the solid disk with or without a mesh, and the perforated disk with a mesh (as showed Fig. S3).

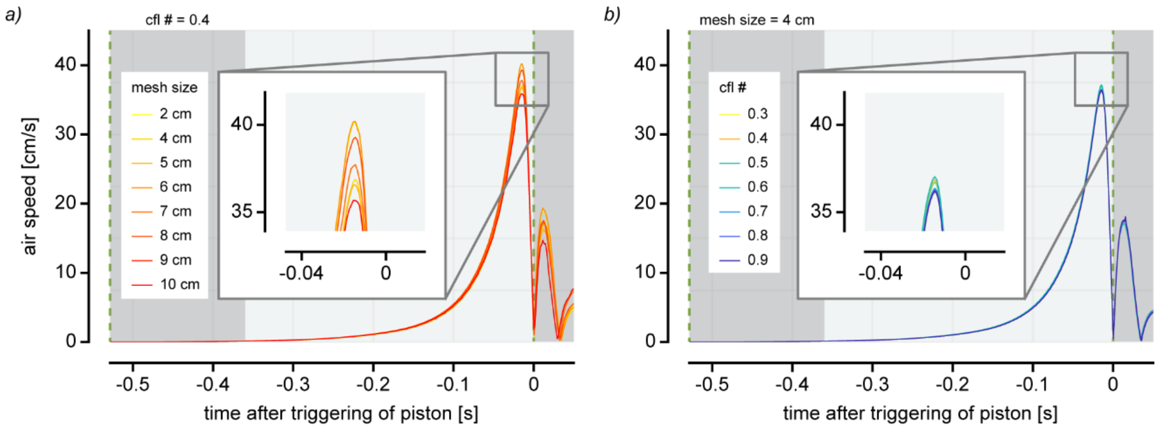


**Fig. S9. Mesh and CFL study.** (*a*,*b*) Comparison between airflow speed from CFD simulations with various mesh sizes or various cfl (Courant–Friedrichs–Lewy) numbers at the same probe position as in supplementary Fig. S8*a*).

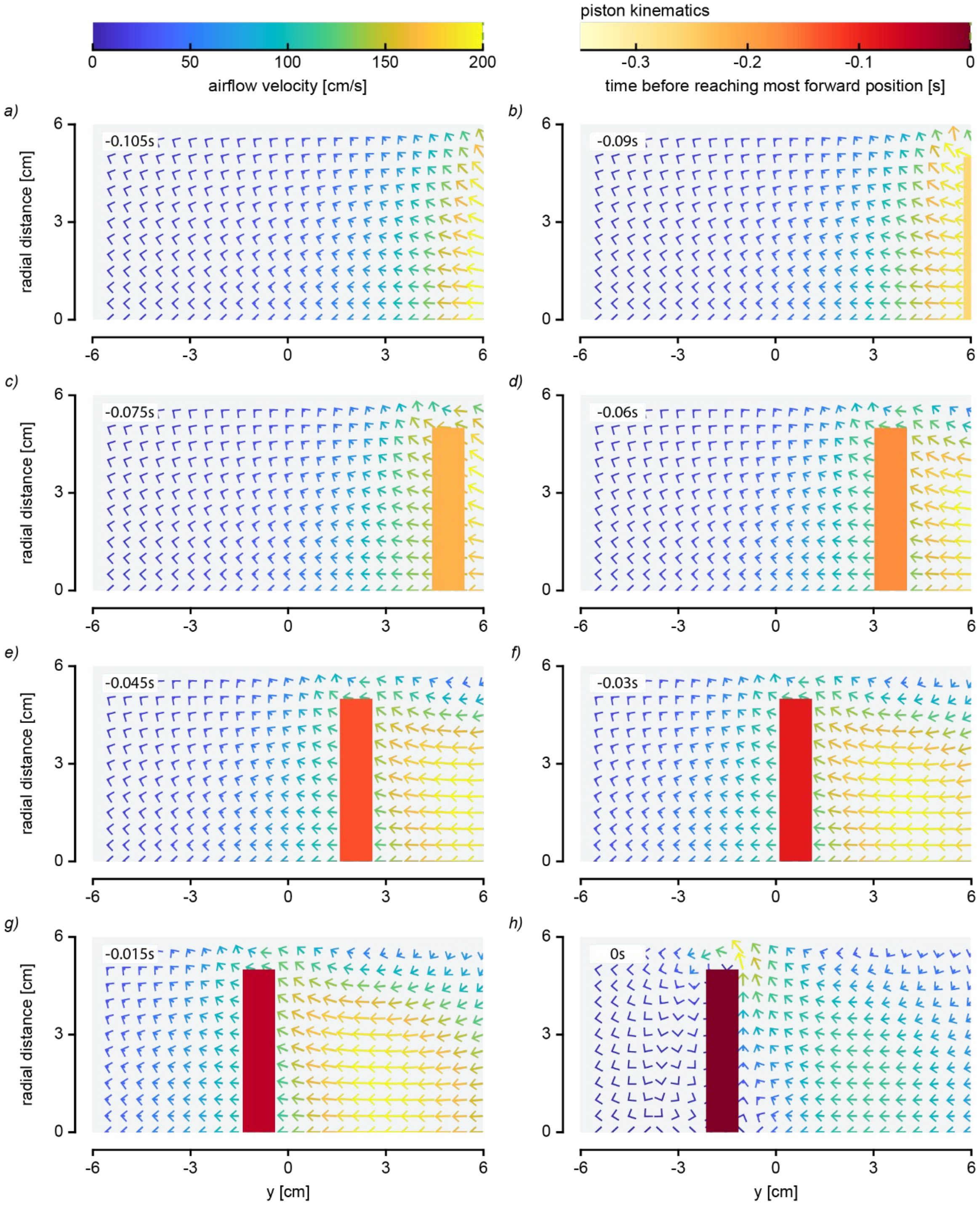


**Fig. S10. Simulated airflow velocity field.** (*a*-*h*) Airflow velocity field from the CFD simulation used in the study. Because the results are following the axial symmetry of the swatter (around y axis), the three-dimensional airflow can be projected into two-dimensional planes. The air is both pushed and pulled by the swatter toward the left, where a mosquito was predicted to be.

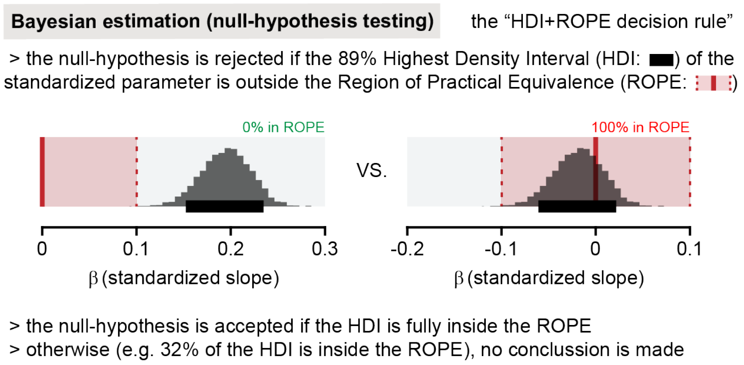


**Fig. S11. Null hypothesis testing with Bayesian estimation.** Examples of two distributions of the estimated mean of a standardized parameter *β* in black. The null hypothesis is rejected (left) if the Highest Density Interval (HDI) is outside the Region Of Practical Equivalence (ROPE). The null-hypothesis is accepted (right) if the full HDI is inside the ROPE.

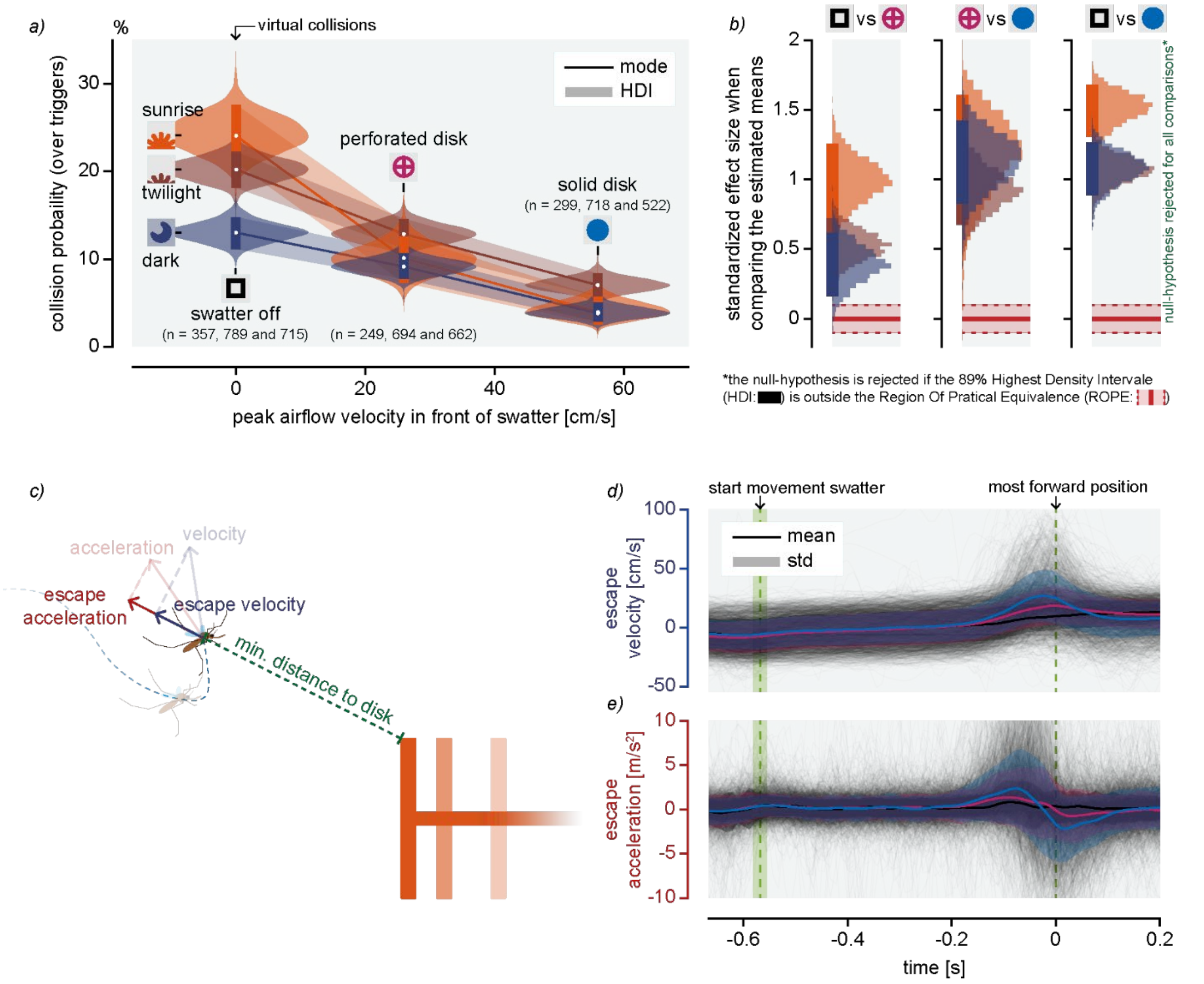


**Fig. S12. Mosquito collision probability and escape dynamics.** (*a*) Distributions of the estimated (using Bayesian estimations) means of mosquito collision probability in the various experimental conditions. The mean collision probability of mosquitoes was higher when the swatter was turned off (virtual collisions) than when the mosquitoes were attacked by the hollow disk or full disk. For all light conditions, mosquito collision probabilities decreased with the increasing peak airflow velocities generated by the different swatter (virtual or not). (*b*) Standardized effect sizes were computed to compare the estimated means of panel *a* (and Fig. 1E). Here, all standardized effect sizes differ significantly from zero. Detailed explanation about null-hypothesis testing is provided in supplementary Fig. S11. (*c*) Schematic showing how the instantaneous escape velocity and acceleration of a mosquito is defined as a function of its relative position with the swatter. (*d*,*e*) Mosquitoes escape velocities and accelerations over time (excluding tracks that resulted in collisions). Mosquitoes are accelerating more and are flying faster when escaping from the solid swatter than from the perforated swatter.

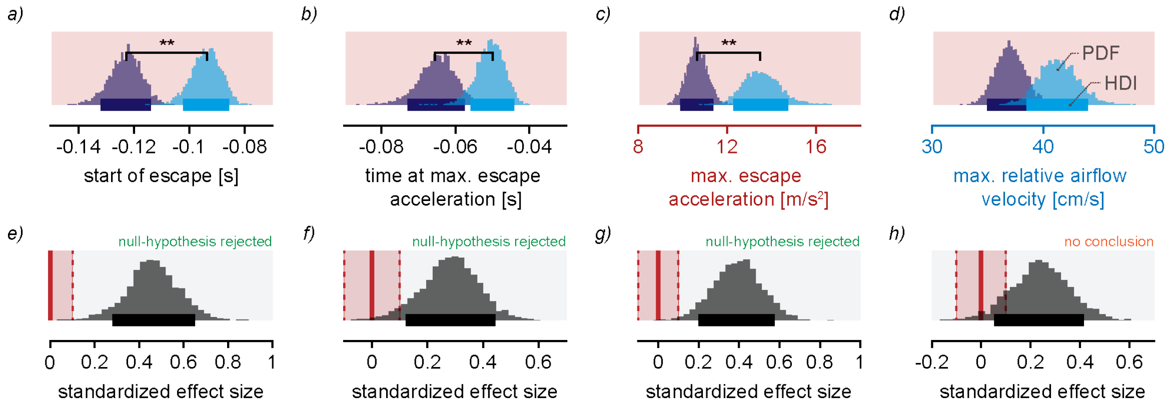


**Fig. S13. Performance metrics of escaping mosquitoes from opaque or transparent disk.** (*a*-*b*) Bayesian estimates of various escape performance metrics: (*a*) Time of entered the escape state; (*b*) time of reached their maximum escape acceleration; (*c*) maximum escape acceleration; (*d*) maximum relative airflow velocity. ** show statistically significant differences; HDI: 89% Highest Density Interval; PDF: Probability Density Function. (*e*-*h*) Standardized effect size of the comparisons between the estimated means of Fig.3I-L. Here the null-hypothesis is rejected for all comparisons except the last, where no conclusion could be made about if the maximum airflow velocity from mosquitoes reference frame differed between the two disk types.

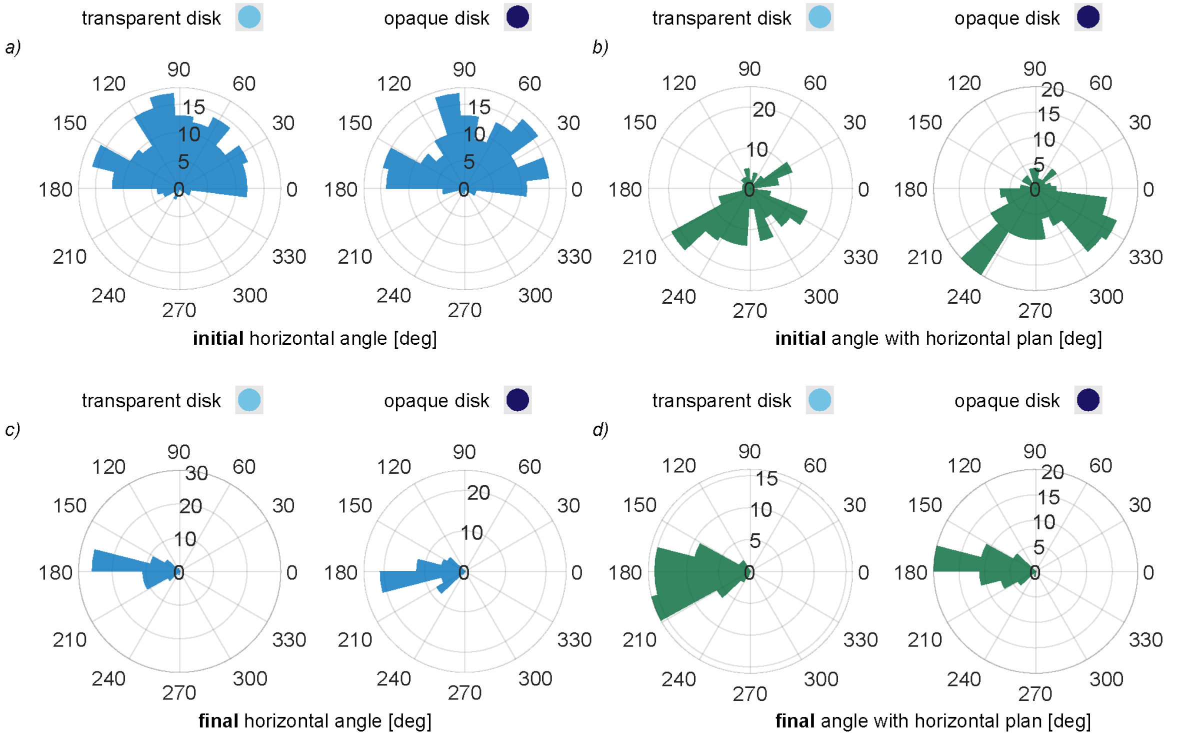


**Fig. S14. Initial and final flight angles.** (*a*,*c*) Rose plots of mosquito flight angle in the horizontal plan at trigger time ((*a*) - initial) and when the swatter reach its most forward position ((*c*) - final). The large majority of tracks are shown to be initially coming from one side because the tracks that had negative initial heading were mirrored. (*b*,*d*) Rose plots of the initial and final mosquito flight angle with the horizontal plan. The swatter is coming from the right (0° angle). Radial scale showing the number of tracks in each bar.

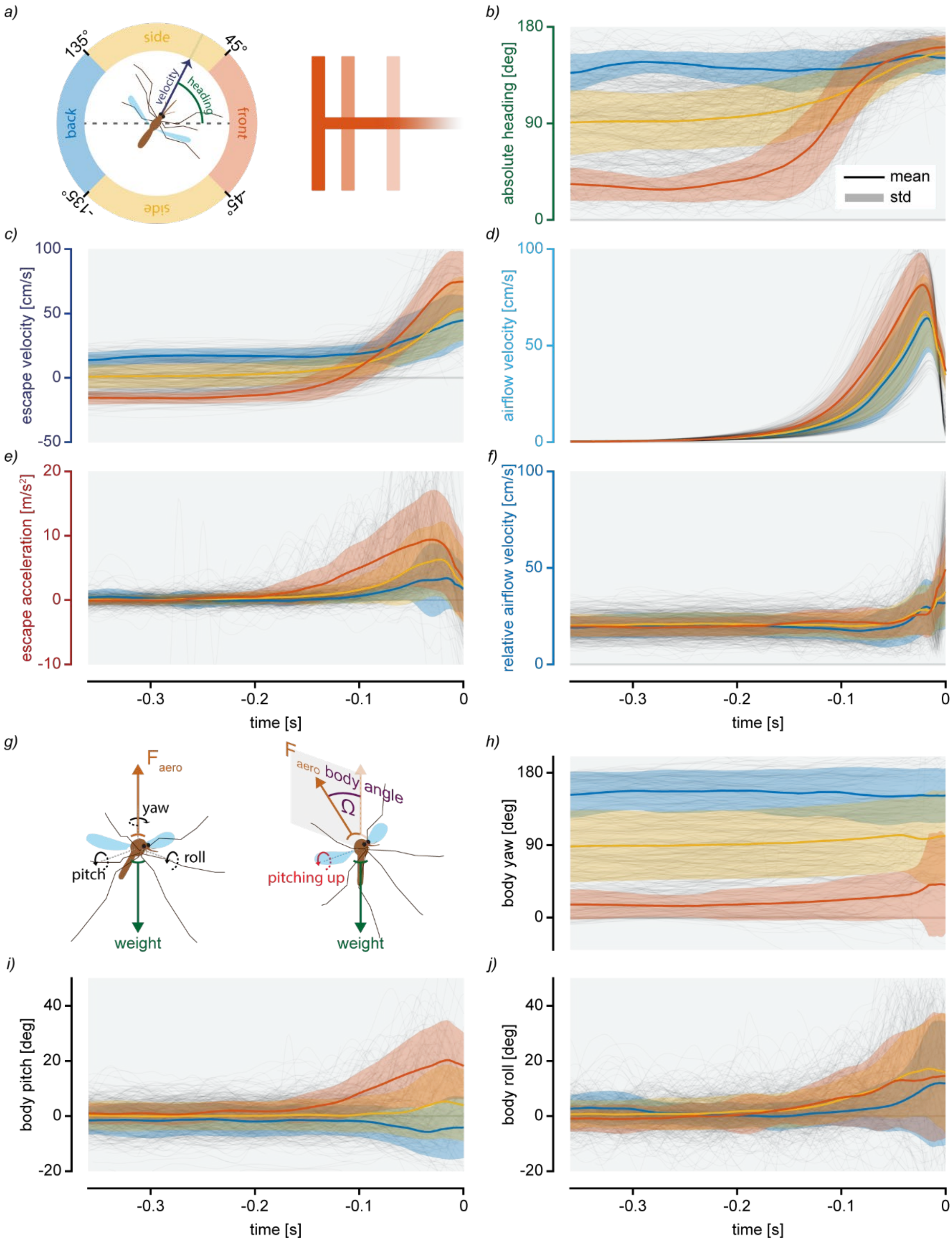


**Fig. S15. Effect of initial heading.** (*a*) Mosquito initial heading is defined as the mean angle between mosquito velocity vector and the direction of the swatter between -0.32 and -0.24 seconds before the swatter reaches its most forward position. (*b-f*) Dynamics of various metrics as a function of mosquito initial heading while escaping. (*g*) Schematic showing how the body yaw, pitch and roll are defined (Tait-Bryan convention). As an example, if a mosquito is pitching up, the force generated by its wings will be redirected backward. Mosquito body tilt angle Ω is defined as the angle between the vertical axis and vector normal to the body of the mosquito (i.e. this vector is directed upward when the mosquito is hovering). When hovering, Ω is equal to zero. (*h-j*) Dynamics of mosquitoes' body tilt angle in function of their initial heading. Around half of mosquito manoeuvres have been mirrored in order for their roll angle to be positive when they rolled away from the swatter (i.e. as if mosquitoes are all flying from the same side). These results confirm that mosquitoes use the so-called "helicopter model" when escaping, as they re-orient the vector of the aerodynamic force generated by their wing away from the danger. For example, mosquitoes pitched up when facing the swatter and pitched down when being attack in the back. They also exhibit higher escape velocities and accelerations when initially facing the swatter than when initially flying away from it.

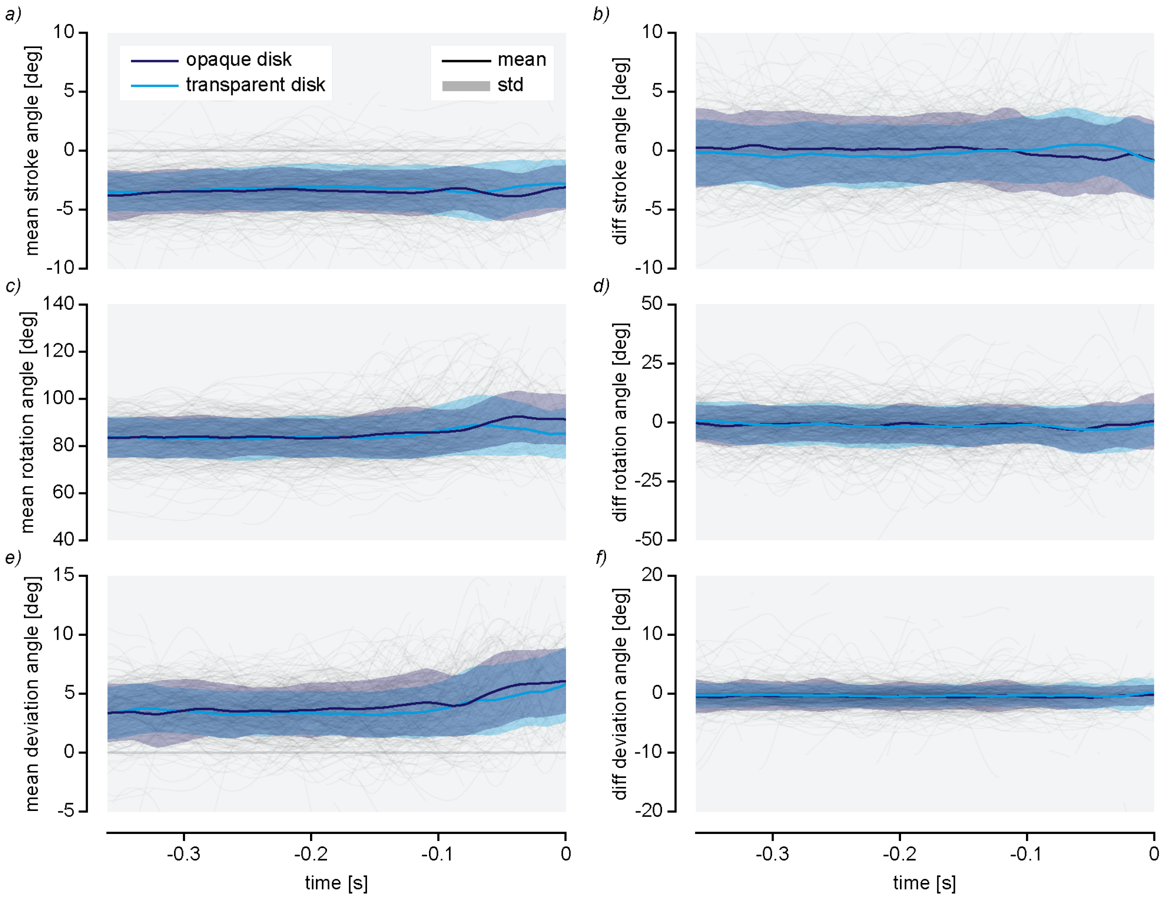


**Fig. S16. Wing angles.** (*a*,*c*,*e*) wingbeat average of the mean both wings of the three wing angles during all the manoeuvres: stroke angle, rotation angle and deviation angle. (*b*,*d*,*f*) wingbeat average of the difference between the right and the left wings of the three wing angles during all the manoeuvres.

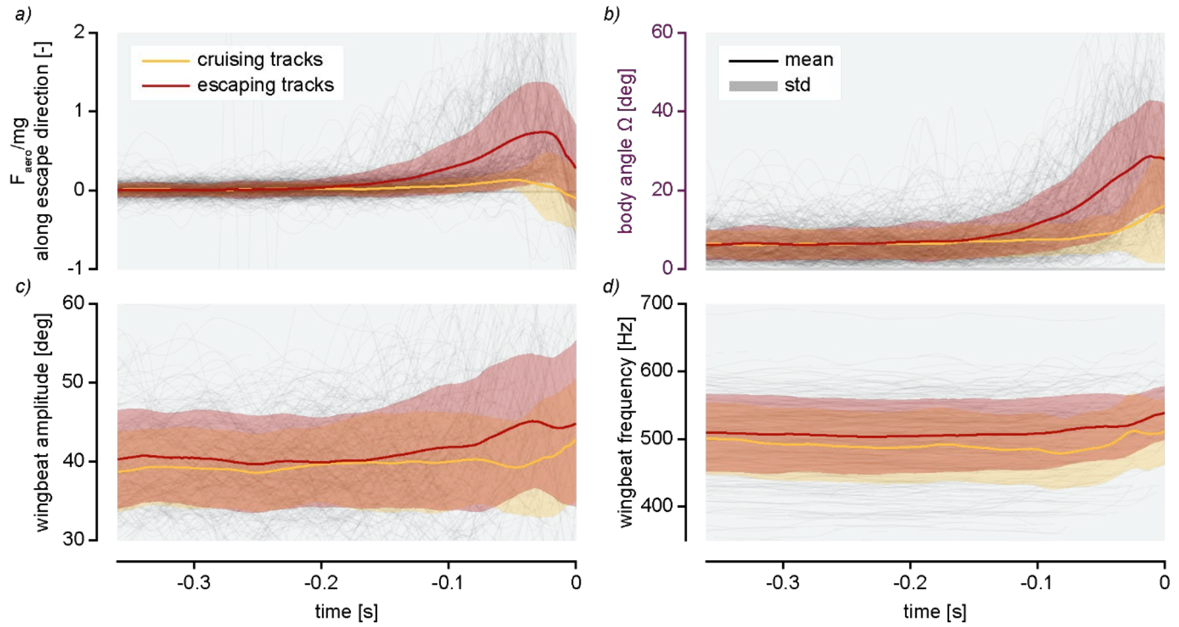


**Fig. S17. Mosquitos actively contribute to their escape maneuvers.** (*a-d*) The increase in observed escape force is correlated with changes in mosquitoes' body tilt angles as well as with increases of their wingbeat amplitudes and frequencies.

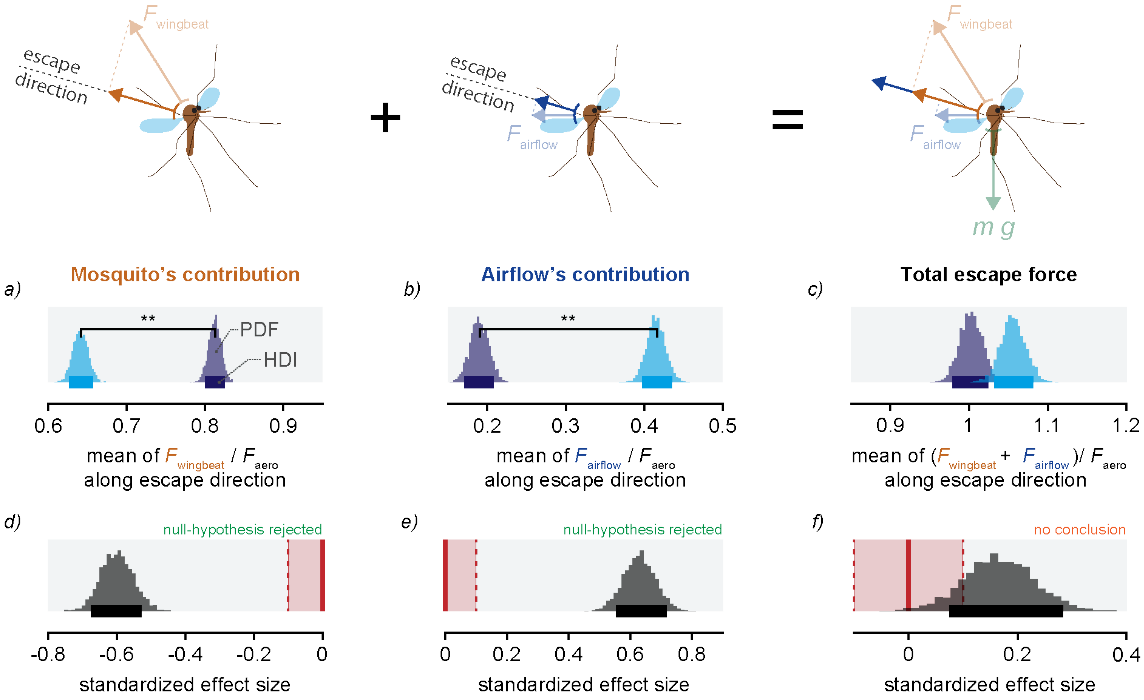


**Fig. S18. Proportion of forces produced during the escapes.** (*a-c*) Distribution of the estimated means of the force proportions (e.g. slopes from Fig. 4D-F)). ** show statistically significant differences; HDI: 89% Highest Density Interval; PDF: Probability Density Function. (*d-f*) Effect size of the comparisons between the force proportions for the transparent and opaque disks. $\vec{F}_{\text{mosquito}}$ was found to contribute significantly more to the total $\vec{F}_{\text{aero}}$with the opaque disk than with the transparent disk.

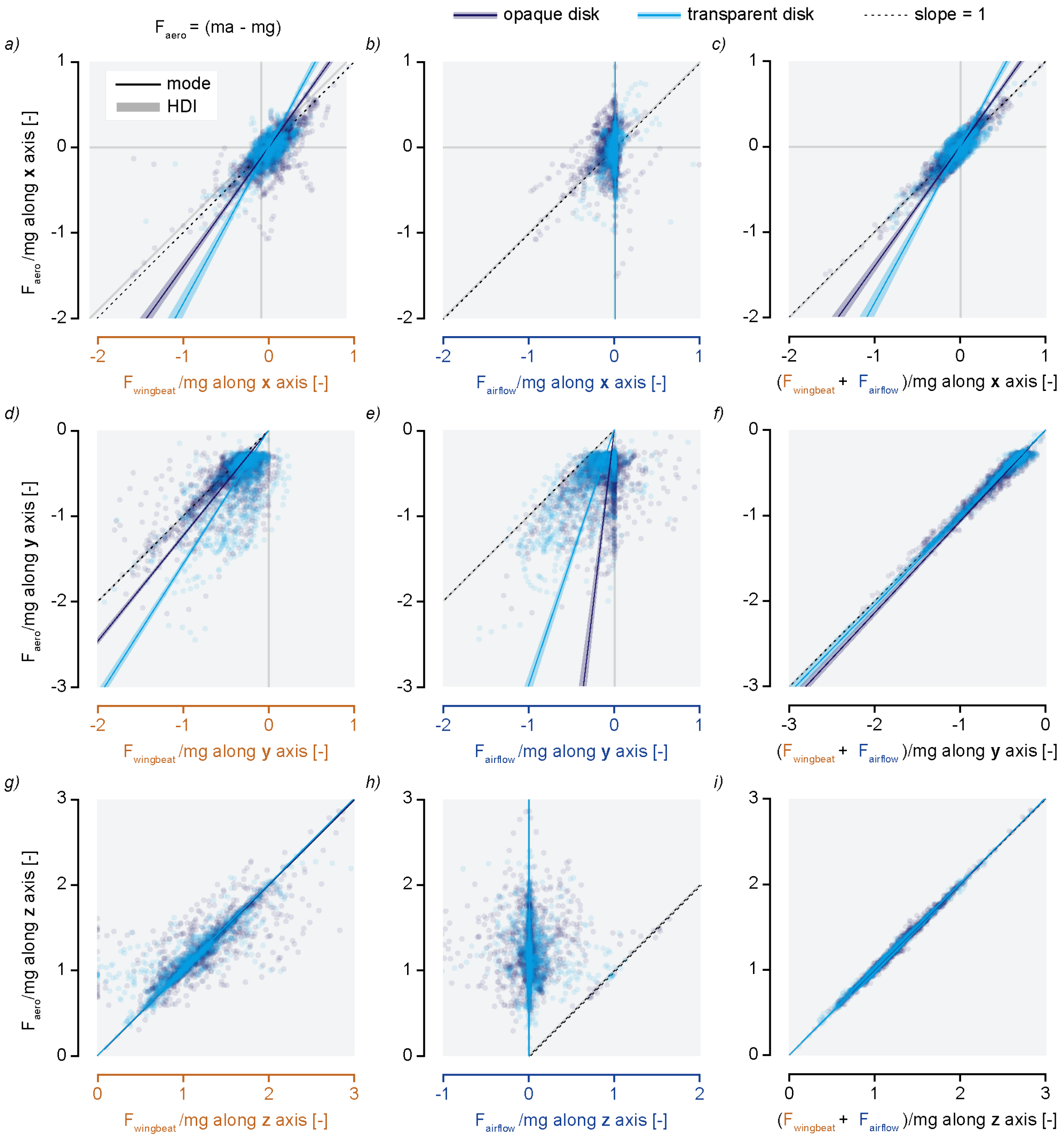


**Fig. S19. Proportion of $\vec{F}_{aero}$ explained by $\vec{F}_{wingbeat}$or $\vec{F}_{airflow}$.** (*a-i*) $\vec{F}_{\mathrm{aero}}$ ($= m\cdot\vec{a} - m\cdot\vec{g}$) in function of estimated $\vec{F}_{\mathrm{mosquito}}$ or $\vec{F}_{\mathrm{airflow}}$ for the transparent and opaque disks. Linear fits have been estimated using Bayesian estimations of the means of the proportion between $\vec{F}_{aero}$ and the corresponding forces in ordinate.

**Legends of supplementary materials not included in this pdf:**

**Movie S1.**

An example of an escape manoeuvre of *Anopheles coluzzii* female mosquito attacked by the mechanical swatter with an transparent disk. The recordings have been slowed down 25 times. The same mosquito is visible on the side and top view (respectively on the left and right of the movie). In the second half of the movie, the airflow velocity field (in light blue) and the acceleration vector (in red) of the mosquito have been added to the recordings.

**Movie S2.**

An example of an escape manoeuvre of *Anopheles coluzzii* female mosquito attacked by the mechanical swatter with an opaque disk. The recordings have been slowed down 25 times. The same mosquito is visible on the side and top view (respectively on the left and right of the movie). In the second half of the movie, the airflow velocity field (in light blue) and the acceleration vector (in red) of the mosquito have been added to the recordings.

**Movie S3.**

An example of an escape manoeuvre of *Anopheles coluzzii* female mosquito attacked by the mechanical swatter with an opaque disk. In this example, the mosquito accelerate away from the swatter twice during the manoeuvre (i.e. two acceleration peaks). The recordings have been slowed down 25 times. The same mosquito is visible on the side and top view (respectively on the left and right of the movie). In the second half of the movie, the airflow velocity field (in light blue) and the acceleration vector (in red) of the mosquito have been added to the recordings.

**Movie S4.**

Results from the tracker used to estimate body and wing kinematics of escaping mosquitoes. The movie shows three views of the same flying mosquitoes over which was projected in two dimensions the three-dimensional skeleton that was fit to the mosquito's body and wings (see Fig. S5 and S6).

**Data S1. (separate file)**

Database of the experiment #1. Flight tracks of all *Anopheles coluzzii* mosquitoes attacked by the (real and virtual) mechanical swatter, and flying in various light conditions. A Matlab .mat file containing three-dimensional tracks of all flying mosquitoes, described as the time t (s) and the spatial {x, y, z} (m) coordinates of the mosquito at each video frame. The coordinates are in meters, and in the world reference frame as defined in figure 1, with z oriented vertically up, and the origin of the coordinate frame at the centre of the flight arena. The trajectories were determined as described in the materials and methods. Metadata for each track includes age, swatter type (hollow or solid) and mode (on or off), light condition, as well as temperature and humidity over time in the arena during each experiment. In addition, the file contains the kinematic of the mechanical swatter, as well as the geometry of the flight arena and the swatter.

**Data S2. (separate file)**

Database of the experiment #2. Flight kinematics of escaping *Anopheles coluzzii* mosquitoes attacked by the mechanical swatter (opaque or transparent disk) in twilight light condition. A Matlab .mat file containing three-dimensional tracks of all escaping mosquitoes as well as all body and wing kinematics parameters (positions and angles) over time. These kinematics parameters were determined using the custom-made three-dimensional tracker (Code S3). Metadata for each track includes age, swatter type (opaque or transparent), as well as temperature and humidity over time in the arena during each experiment. In addition, the file contains the kinematic of the mechanical swatter, as well as the geometry of the flight arena and the swatter.

**Data S3. (separate file)**

Computation fluid dynamic (CFD) results. Means and standard deviation of the airflow velocities induced by the swatter movement can be found in .csv files (one for each time step (1 ms) and in polar coordinates (r,theta,y)). The piston kinematic and the probe positions can also be found in separated .csv files.

**Code S1. (separate file)**

Python code used to track mosquitoes body and wings kinematics. Recordings of mosquitoes were pre-processed (cropped and stitched) to be analysed with Deeplabcut. Then a three-dimensional mosquito skeleton was fitted to the two-dimensional tracking results from Deeplabcut to estimate the body and wing kinematics. Scripts to generate stroboscopic images and example videos are also included.

**Code S2. (separate file)**

Analysis codes. Contains all original Matlab codes that were written to perform the analysis of the article and to generate the panels of Fig. 1-4. Also contains JAGS modelling codes used for doing the Bayesian estimations in Fig. 1, 3-4. Instructions to run the full analysis can be found in the readme.md.
